# Peroxiredoxin 2 Regulates DAF-16/FOXO Mediated Mitochondrial Remodelling in Response to Exercise that is Disrupted in Ageing

**DOI:** 10.1101/2024.05.13.593975

**Authors:** Qin Xia, Penglin Li, José C. Casas-Martinez, Antonio Miranda-Vizuete, Emma McDermott, Peter Dockery, Katarzyna Goljanek-Whysall, Brian McDonagh

**Affiliations:** Discipline of Physiology, School of Medicine; Apoptosis Research Centre, University of Galway, Ireland; Instituto de Biomedicina de Sevilla, Hospital Universitario Virgen del Rocío/CSIC/Universidad de Sevilla, Sevilla, Spain; Centre for Microscopy and Imaging, Discipline of Anatomy, School of Medicine, University of Galway, Ireland; Institute of Lifecourse and Medical Sciences, University of Liverpool, UK

**Keywords:** Ageing, Peroxiredoxin 2, Exercise, *C. elegans*, Mitochondrial ER Contact Sites, DAF-16

## Abstract

Ageing is associated with mitochondrial dysfunction and increased oxidative stress. Exercise generates endogenous reactive oxygen species (ROS) and promotes rapid mitochondrial remodelling. We investigated the role of Peroxiredoxin 2 (PRDX-2) in mitochondrial adaptations to exercise and ageing using *Caenorhabditis elegans* as a model system. PRDX-2 was required for the mitochondrial remodelling in response to exercise mediated by DAF-16 nuclear localisation. Employing an acute exercise and recovery cycle, we demonstrated exercise-induced mitochondrial ER contact sites (MERCS) assembly and mitochondrial remodelling dependent on PRDX-2 and DAF-16 signalling. There was increased mitochondrial fragmentation, elevated ROS and an altered redox state of PRDX-2, concomitant with impaired DAF-16 nuclear localisation during ageing. Similarly, the *prdx-2* mutant strain exhibited increased mitochondrial fragmentation and a failure to activate DAF-16 required for mitochondrial fusion. Collectively, our data highlight the critical role of PRDX-2 in orchestrating mitochondrial remodelling in response to a physiological stress by regulating DAF-16 nuclear localisation.

**Graphical abstract:** 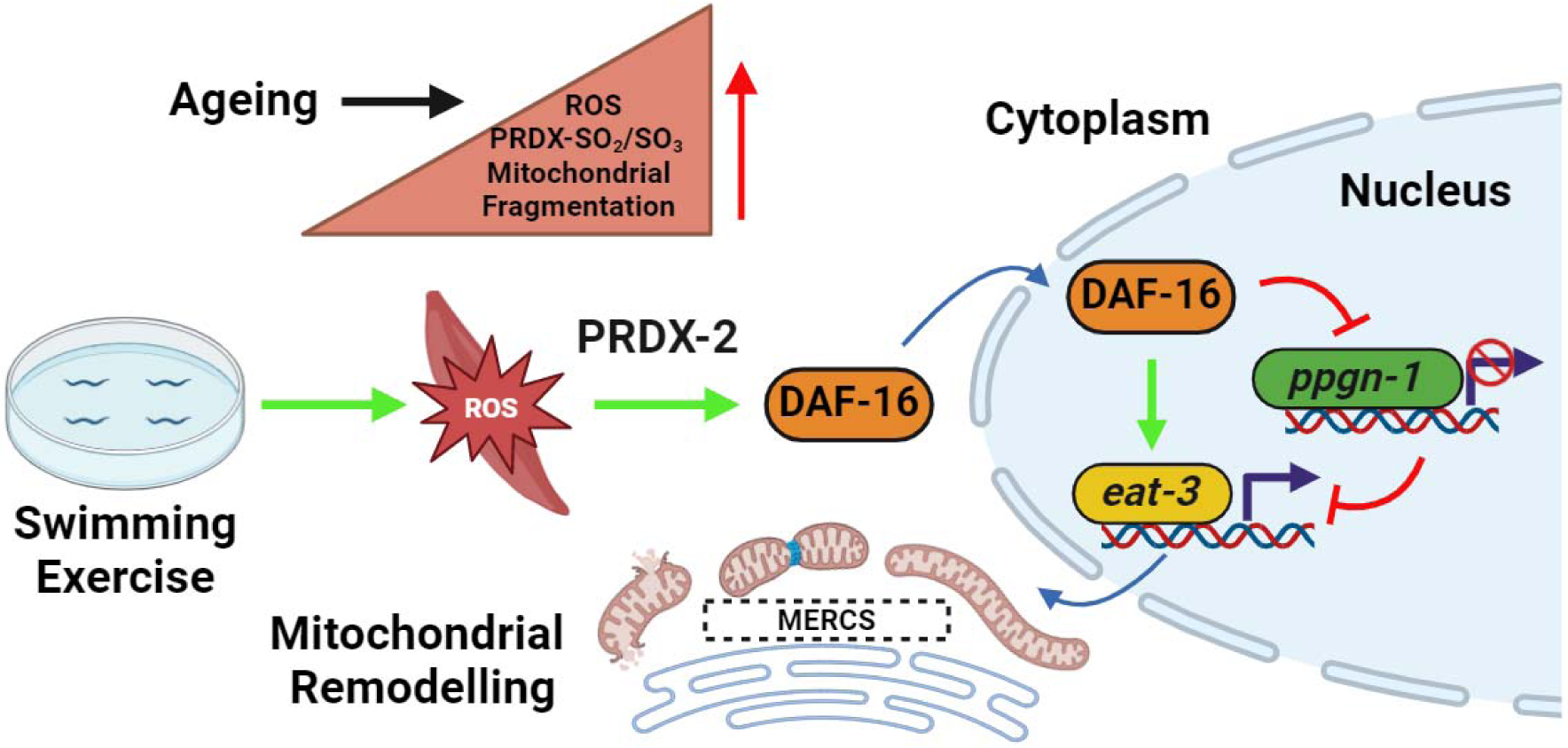

**Highlights:**

- Exercise generates ROS and promotes mitochondrial remodelling dependent on DAF-16.
- Exercise induces mitochondrial ER contact site assembly and mitochondrial dynamics.
- Ageing and loss of PRDX-2 results in disrupted mitochondrial fusion.
- The redox state of PRDX-2 determines appropriate DAF-16 nuclear localisation.

## 1. Introduction

Ageing results in a gradual decline in physiological function, leading to compromised overall health and increased susceptibility to diseases and mortality [1]. Age-related disorders, such as diabetes, sarcopenia and neurodegenerative conditions, pose a significant and growing public health challenge [2, 3]. Exercise is recognised as a nonpharmacological intervention to mitigate the adverse effects of ageing. Physical exercise is essential for preserving and enhancing skeletal muscle function, ameliorating age-related muscle atrophy and providing systemic benefits against various age-related diseases [4, 5]. During exercise, skeletal muscle mitochondria undergo significant structural remodelling to meet energy and nutritional requirements, impacting cellular metabolism and signalling [6, 7]. The decline in mitochondrial function is a hallmark of ageing [8] and the accumulation of dysfunctional mitochondria with age has been observed in *C. elegans* [9]. Damaged or dysfunctional components of mitochondria can be selectively removed via fission by the transfer to the lysosome for degradation or mitophagy [10]. Another critical mechanism for mitochondrial health is fusion, where individual mitochondria merge to form a singular, enlarged structure [11]. The maintenance of mitochondrial quality, encompassing processes such as mitochondrial biogenesis, fission and fusion, is critically important in governing both the structure and functionality of mitochondria. In *C. elegans*, mitochondrial fission involves DRP-1 (DRP1 in mammals), while fusion requires FZO-1 (MFN1/2 in mammals) for outer membrane fusion and EAT-3 (OPA1 in mammals) for inner membrane fusion [12]. Overexpression of DRP-1 leads to mitochondrial hyper-fragmentation, whereas suppression of DRP-1 results in fragmented mitochondrial matrices but fused outer mitochondrial membranes [13]. Mitochondrial fusion mutant strains (*eat-3* and *fzo-1*) also display increased mitochondrial fragmentation [14].

The Forkhead Box O (FOXO) family of transcription factors play crucial roles in regulating diverse cellular processes such as oxidative stress response, mitochondrial function and cellular metabolism [15]. In *C. elegans*, the FOXO ortholog, DAF-16, exhibits heightened nuclear localisation in response to elevated reactive oxygen species (ROS) and following a swimming exercise [16, 17]. Moreover, DAF-16 has been reported to regulate mitochondrial morphology [18]. The mitochondrial proteases SPG-7 and PPGN-1 target the mitochondrial fusion protein EAT-3, DAF-16 nuclear localisation results in transcriptional repression of *spg-7* and *ppgn-1*, allowing the negative regulation of EAT-3 to be alleviated and thereby promoting mitochondrial fusion [18].

Mitochondrial morphology is also thought to determine fuel substrate use, whereby an increase in mitochondrial fragmentation increases fatty acid oxidation [19]. It was demonstrated that fragmented mitochondria promote Carnitine O-palmitoyltransferase 1 (CPT1) regulated long chain fatty acid oxidation [19]. Our previous proteomic analysis following a swim exercise in wild type (N2) *C. elegans* demonstrated an increased abundance of proteins involved in fatty acids oxidation and decreased abundance of fatty acid anabolism and the most upregulated protein was CPT-1 [20]. These results indicated that under conditions of bioenergetic stress such as during exercise, there was a promotion of mitochondrial fatty acid oxidation and increased mitochondrial fragmentation.

Mitochondria and endoplasmic reticulum (ER) are key regulatory hubs in maintaining cellular homeostasis and have a synergistic relationship that determines their function and adaptability to the cellular environment. Communication between these organelles is mediated by Mitochondrial ER contact sites (MERCS), allowing the exchange of metabolites, lipids and calcium [21, 22]. MERCS are dynamic and relatively stable structures between mitochondria and ER (< 50 nm) that remodel in response to cellular signalling events, that can affect the function of both organelles. Recent evidence would suggest that during acute ER stress, there is activation of the adaptive Unfolded Protein Response (UPR), resulting in ER and MERCS remodelling [23]. MERCS have been identified as regulating the sites of mitochondrial fission and there is some evidence to suggest they also play a role in mitochondrial fusion [22, 24]. During fission, the ER constricts mitochondria at the site of fission using Drp1 [25]. Furthermore, the sites of mitochondrial fission have distinct ROS signatures that may regulate formation of MERCS, fission at the periphery or tip results in mitochondrial fragments destined for degradation while midzone fission is preferential for mitochondrial dynamics [26]. Disruption of mitochondrial dynamics has been reported in a wide variety of age-related diseases including neurodegeneration and sarcopenia, associated with an accumulation of dysfunctional mitochondria [27]. An increase in MERCS formation has been proposed to be involved in cell senescence and in models of neurodegenerative disease [28, 29]. In skeletal muscle, sarcomeres are surrounded by mitochondria and sarco/endoplasmic reticulum, essential for Ca^2+^ regulation. However, decreased MERCS formation has been reported with age [30]. Energy stress and subsequent AMPK activation has been demonstrated in cell models to promote autophagy and MERCS formation [31]. Introducing an exercise protocol that promotes a mild ER stress response and induces mitochondrial remodelling, will ultimately result in an improved bioenergetic profile. During exercise, there is a site-specific elevation of endogenous ROS in skeletal muscle [32, 33]. However, it has been reported in a number of different studies that there is a chronic basal elevation of ROS in muscle with age [34, 35]. The acute elevation in ROS following a physiological stress such as exercise activates specific signalling pathways including NRF2 and NF-κB [36, 37], promoting a beneficial adaptive response. However, the majority of prior investigations mainly focused on static redox states, with limited exploration of dynamic responses in both young and elderly individuals to redox stress. The Peroxiredoxins (PRDXs) family, constitute up to 1% of cellular protein content and are generally considered as a group of antioxidant enzymes with peroxidase activity [38]. Peroxiredoxin 2 (PRDX-2) functions as a peroxidase to mitigate ROS during stress and has been demonstrated to be required for the beneficial adaptation to exercise in *C. elegans* [20]. Understanding the specific role of PRDX-2 in regulating mitochondrial dynamics during exercise and ageing is essential.

In this study, the nematode *C. elegans* was utilised as a model system to investigate the intricate interplay between ageing and exercise, with a specific focus on elucidating the role of PRDX-2 in mitochondrial dynamics. We demonstrated that exercise promoted mitochondrial dynamics, MERCS formation and fatigue in *C. elegans*. A 24-hour recovery period was adequate to increase mitochondrial fusion in young worms. Old worms exhibited fragmented mitochondria, no or delayed recovery response, an accumulation of basal ROS, an increase in hyperoxidised Peroxiredoxins and decreased fitness. The *prdx-2* mutant strain had increased mitochondrial fragmentation but did not activate mitophagy and displayed reduced physical fitness. Additionally, the acute swim exercise induced nuclear localisation of DAF-16 in the adult N2 strain promoting mitochondrial fusion. However, this response was not observed in the ageing N2 strain or the *prdx-2* mutant strain, which did not stimulate mitochondrial fusion as a result of a failure to stimulate DAF-16 dependent signalling. Our data results demonstrate blunted mitochondrial remodelling and an accumulation of ROS with age associated with disruption in the redox state of PRDX-2, highlighting the crucial role of PRDX-2 in coordinating mitochondrial adaptation in response to exercise and ageing.

## 2 Results

### 2.1 Redox mediated adaptation to exercise diminishes with age through a PRDX-2 dependent pathway

To explore the adaptive response to ageing and exercise, we employed *C. elegans*, a model organism that has been validated as an effective platform for studying physiological responses to exercise [20, 39, 40]. A 90-minute swimming exercise has been demonstrated to elicit muscle fat consumption, increase locomotory fatigue and elevate mitochondrial ROS levels [41]. Therefore, we conducted a single 90-minute swim exercise session at distinct developmental stages, adult (D4), late middle age (D8) and old (D12) worms (Figure 1A) in N2 wild type strain. Initially, worms were subjected to diverse forms of oxidative stress, including exposure to paraquat (PQ) and arsenite (AS) following an exercise regimen and subsequent recovery. A discernible reduction in the adaptive response capacity was observed with progressive ageing. The resistance to both PQ and AS decreased immediately after acute exercise (Figures 1B and 1C). Subsequently, a recovery was observed 24 hours post-exercise in the younger D4 and D8 worms, contrasting with the absence of a recovery in D12 worms (Figures S1A-S1H). The mean lifespan of each condition is detailed in Table S1. These results demonstrate a decreased capacity to withstand stress with age and older worms are not able to recover following a physiological exercise stress.

**Figure 1.**
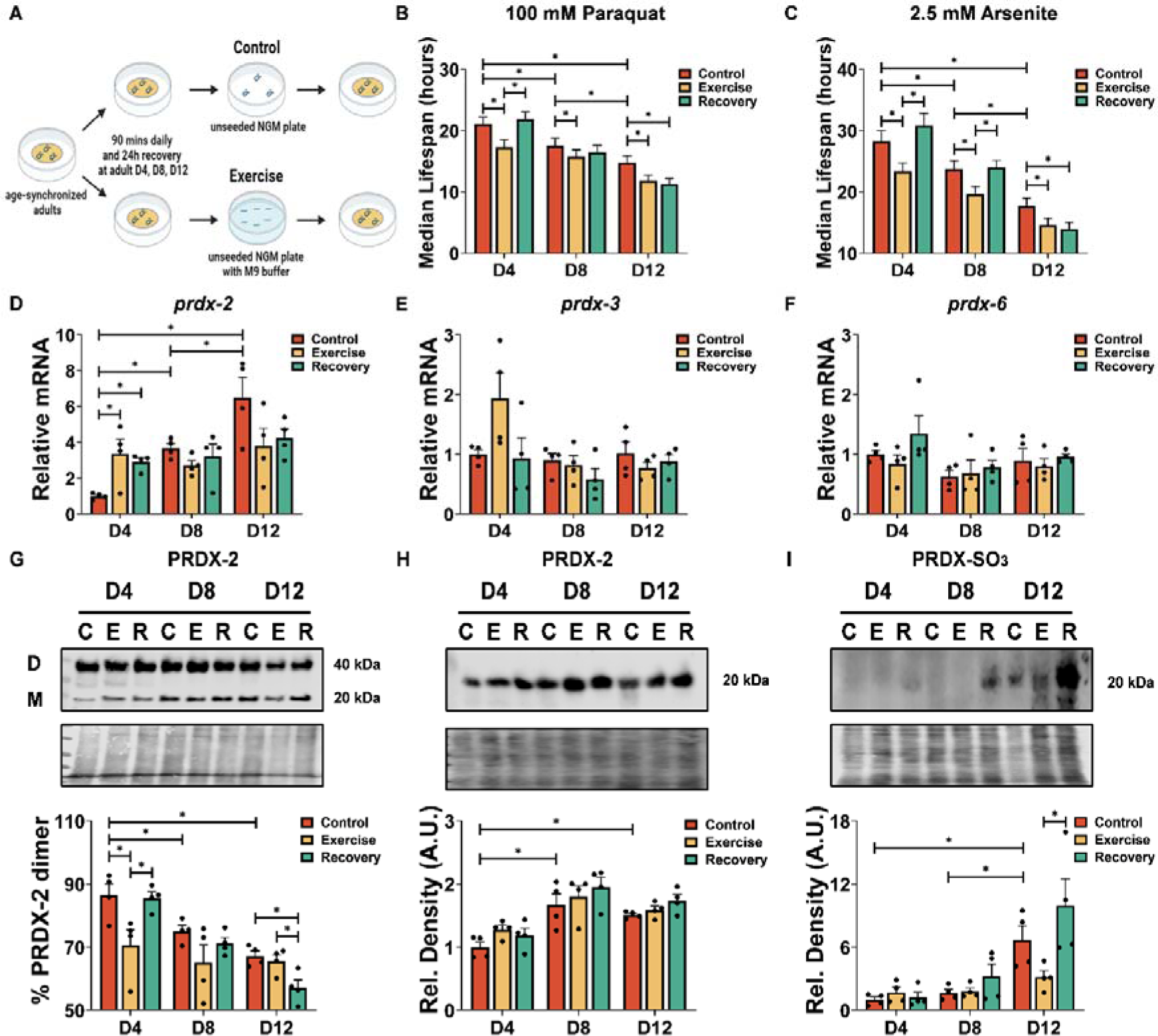
Decreased survival with exposure to oxidative stress, alterations in Peroxiredoxin mRNA and PRDX-2 in response to ageing and exercise. (A) Schematic of acute swimming protocol for *C. elegans* at different ages. (B and C) Survival to paraquat and arsenite following acute exercise as well as ageing (n = 60). (D-F) mRNA levels of *prdx-2*, *prdx-3* and *prdx-6* following acute exercise and ageing (n = 4). (G-I) Percentage dimer/ total (dimer+monomer) ratio of PRDX-2 (G), protein levels of PRDX-2 (H) and PRDX-SO_2_/SO_3_ (I) (n = 4). Graphs are the normalised relative means ± SEM and *p*-value of < 0.05 was considered as statistically significant *(*p* < 0.05). (B: D4: C vs E = 0.0034, E vs R = 0.001; D8: C vs E = 0.0435; D12: C vs E = 0.022, C vs R = 0.0076; D4 vs D8: = 0.0266; D4 vs D12 < 0.0001; D8 vs D12 = 0.0181. C: D4: C vs E = 0.0114, E vs R = 0.0009; D8: C vs E = 0.0143, E vs R = 0.0079; D12: C vs E = 0.031; C vs R = 0.0048; D4 vs D8: = 0.0136; D4 vs D12 < 0.0001; D8 vs D12 = 0.0023. D: D4: C vs E = 0.017, C vs R = 0.0485; D4 vs D8 = 0.0445; D4 vs D12 = 0.0006. G: D4: C vs E = 0.0325; E vs R = 0.0432; D12: C vs R = 0.0173, E vs R = 0.0402; D4 vs D8 = 0.0237; D4 vs D12 = 0.0009. H: D4 vs D8 = 0.0063; D4 vs D12 = 0.0271. I: D12: E vs R = 0.0475; D4 vs D12 = 0.002; D8 vs D12 = 0.0044)

Given the previously demonstrated involvement of PRDX-2 in redox-mediated adaptation to chronic exercise in our prior investigations [20], we conducted qPCR and Western blot analyses following a regimen of acute exercise and subsequent recovery. Initially, we examined the mRNA expression levels of the Peroxiredoxin genes in *C. elegans*, encompassing *prdx-2*, *prdx-3* and *prdx-6*, in response to the exercise and recovery protocol. Intriguingly, our observations unveiled a distinctive sensitivity pattern, with only *prdx-2* exhibiting an adaptive response to both acute exercise and the ageing processes. Specifically, following acute exercise and post-exercise in D4 worms, *prdx-2* but not *prdx-3* or *prdx-6* demonstrated an elevated expression level (Figures 1D-1F). There was no significant change of *prdx-2* expression in D8 and D12 worms in response to exercise. Furthermore, a gradual increase in the expression of *prdx-2* was observed in control conditions at D8 and D12 during the ageing process (Figure 1D). The ratio of PRDX-2 Dimer:Monomer was analysed using non-reducing gel electrophoresis. Exercise resulted in a pronounced shift in the dimerisation of PRDX-2 in D4 worms but returned to control levels following a recovery period (Figure 1G). D8 and D12 worms had increased expression of PRDX-2 but decreased % PRDX-2 dimer formation compared to D4 worms (Figures 1G and 1H). There was also an increase in hyperoxidised Peroxiredoxins in D12 worms compared to D4 and D8 worms, which increased further following the recovery period (Figure 1I). These findings demonstrate that resistance to oxidative stress declines with ageing and exercise was associated with distinct changes in the sensitivity of the redox state of PRDX-2 to exercise and age.

### 2.2 PRDX-2 is required for mitochondrial adaptation in response to exercise

In order to determine the role of PRDX-2 in mitochondrial adaptation in response to exercise, we performed MitoTracker, MitoSOX and DCFDA staining using N2 and *prdx-2* (gk169) mutant strains throughout a structured exercise and recovery regimen at distinct developmental stages. A progressive reduction in MitoTracker Red staining intensity, indicative of mitochondrial membrane potential, was noted at D8 and D12 worms relative to D4 worms control groups in both N2 and *prdx-2* mutant strains (Figure 2A). Subsequent analysis revealed an immediate increase in staining intensity and mitochondrial membrane potential post-acute swim, with recovery to control levels observed 24 hours after exercise in D4 and D8 worms in the N2 strain. This trend was absent in old D12 wild type N2 strain. MitoTracker red staining decreased in the *prdx-2* mutant strain and unlike the N2 strain, there was no increase in staining at any age following the exercise protocol. Notably, a gradual decline in the recovery rate (Δ24h-0h) was observed with age in both N2 and *prdx-2* mutant strains, with N2 consistently exhibiting a superior recovery rate compared to the *prdx-2* mutant strain across all developmental stages (Figure 2A). To assess the adaptive response to exercise-induced ROS, we utilised DCFDA staining as a proxy for overall cellular ROS and MitoSOX staining as an indicator for mitochondrial ROS. We observed an elevation in DCFDA and MitoSOX intensity at D4 and D8 post-exercise in both N2 and *prdx-2* mutant strains (Figures 2B and S2A). However, this increase was transient in the N2 strain and returned to basal levels following the recovery period but remained elevated in the *prdx-2* mutant strain at D4 and D8. There was a progressive escalation of DCFDA and MitoSOX intensity with advancing age compared to D4 worms in both strains. Remarkably, in old worms at D12, an immediate increase of DCFDA and MitoSOX intensity was not observed directly after exercise, instead a significant increase was noted 24 hours later (Figures 2B and S2A). The *prdx-2* mutant strain was crossed with a reporter for SKN-1 activation, CL2166 *dvIs19[(pAF15)gst-4p::GFP::NLS] III* to assess SKN-1 (ortholog of NRF2) activation during both exercise and ageing process. The staining analysis revealed a gradual elevation of SKN-1 activity at D8 and D12 in both N2 and *prdx-2* mutant strains (Figure S2B). Furthermore, there was an increase in SKN-1 activity 24 hours post-acute exercise at D4, although this effect was not observed at D8 or D12 in the N2 strain. Intriguingly, a decrease in SKN-1 activity was noted during the recovery phase in *prdx-2* mutants at D8 and D12. Consistent with MitoTracker staining results, the SKN-1 recovery rate exhibited a decline with ageing in both N2 and *prdx-2* mutant strains, the N2 strain demonstrated a superior recovery rate compared to the *prdx-2* mutant throughout the lifespan (Figure S2B). Collectively, the data indicate an acute increase in ROS and mitochondrial membrane potential following exercise in the N2 strain that returns to baseline levels following a recovery in D4 and D8 worms. There was a gradual increase in overall ROS with age and a decline in mitochondrial membrane potential. The *prdx-2* mutant strain has reduced mitochondrial membrane potential and exercise induced an increase in ROS at D4 and D8, which did not return to baseline following a recovery period. The results highlight the essential role of PRDX-2 in mediating the adaptive response to exercise.

**Figure 2.**
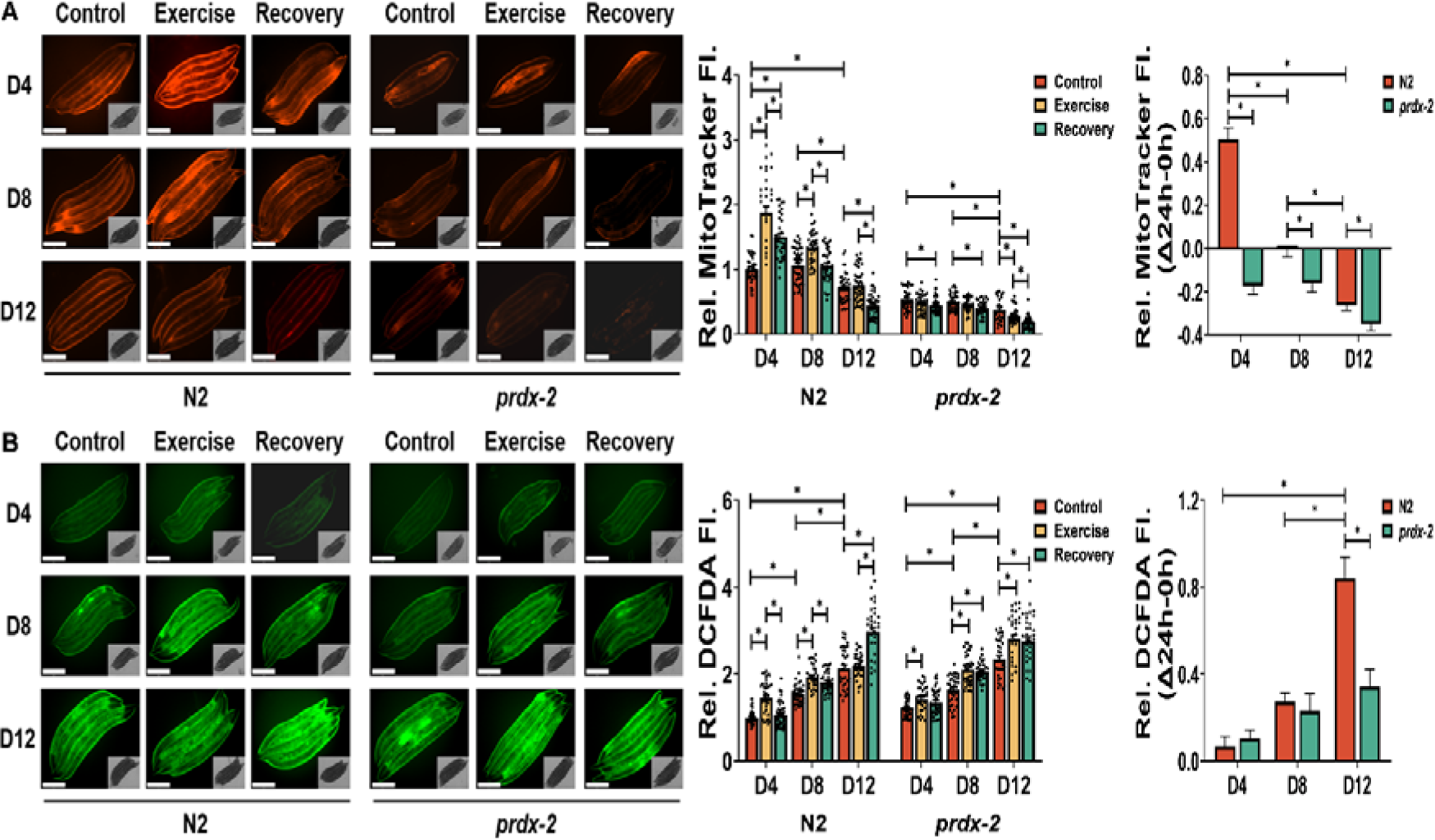
Ageing and loss of PRDX-2 result in decreased membrane potential and increased ROS resulting in disrupted adaptation to exercise. (A-B) MitoTracker red staining as an indicator of mitochondrial membrane potential (A) and DCFDA staining for intracellular ROS (B) following acute exercise at different stages, scale bar = 275 μm, n = 30-40. Graphs are the normalised relative means ± SEM and *p*-value of < 0.05 was considered as statistically significant *(*p* < 0.05). *p* values (A: in N2 worms: D4: C vs E < 0.0001, C vs R < 0.0001, E vs R = 0.0002; D8: C vs E < 0.0001, E vs R < 0.0001; D12: C vs R < 0.0001, E vs R < 0.0001; D4 vs D12 < 0.0001; D8 vs D12 < 0.0001; in *prdx-2* worms: D4: C vs R = 0.0183; D8: C vs R = 0.0079; D12: C vs E = 0.0361, C vs R < 0.0001, T vs R = 0.0086; D4 vs D12 = 0.0001; D8 vs D12 = 0.005; recovery rate: N2: D4 vs D8 = 0.0035; D4 vs D12 = 0.0007; *prdx-2*: D4 vs D12 = 0.002; D8 vs D12 = 0.0022; N2 vs *prdx-2*: D4 < 0.0001; D8 = 0.0067; D12 = 0.0295. B: in N2 worms: D4: C vs R < 0.0001, E vs R < 0.0001; D8: C vs E < 0.0001, C vs R < 0.0001; D12: C vs R < 0.0001; E vs R < 0.0001; D4 vs D8 < 0.0001; D4 vs D12 < 0.0001; D8 vs D12 < 0.0001. in *prdx-2* worms: D4: C vs E = 0.005; D8: C vs E < 0.0001, C vs R < 0.0001; D12: C vs E = 0.001, C vs R = 0.0035; D4 vs D8 < 0.0001; D4 vs D12 < 0.0001; D8 vs D12 < 0.0001. recovery rate: N2: D4 vs D12 < 0.0001; D8 vs D12 < 0.0001; *prdx-2*: D4 vs D12 = 0.0297; N2 vs *prdx-2*: D12 = 0.0002)

### 2.3 Loss of PRDX-2 results in fragmented mitochondria and disrupted mitochondrial dynamics

To elucidate the impact of PRDX-2 on mitochondria in response to exercise, the *prdx-2* mutant strain was crossed with the muscle mitochondrial reporter strain SJ4103 *zcIs14 [myo-3::GFP(mit)]* for monitoring mitochondrial morphology and IR2539 *unc-119(ed3); Ex[pmyo-3TOMM-20::Rosella;unc-119(+)]* to determine mitophagy. Mitochondrial morphology was quantified within the body-wall muscle of *C. elegans* (Figure 3A), a tissue displaying numerous parallels to mammalian skeletal muscle [42]. This analysis entailed the classification of mitochondrial morphology into five distinct categories, each indicative of a stepwise escalation in the levels of mitochondrial fragmentation and disorganisation (Figure 3B). The majority of body wall muscles exhibited plentiful and interconnected mitochondria in D4 N2 worms. At D8 and D12, there was an increase in mitochondrial fragmentation and progressive impairment in mitochondrial connectivity (Figure 3A). Furthermore, acute exercise induced mitochondrial fragmentation in body wall muscles, with subsequent restoration of filamentous mitochondria observed 24 hours post-exercise in D4 and D8 worms, but not in D12 worms (Figure 3B). The *prdx-2* mutant strain exhibits an accelerated ageing phenotype [20, 43] and a highly fragmented mitochondrial network was apparent at D4 that further deteriorates with age, suggesting a critical role for PRDX-2 in preserving mitochondrial integrity during both acute exercise and ageing process. The evaluation of mitochondrial turnover involved the use of a mitophagy reporter strain *[myo-3p tomm-20::Rosella]*. This Rosella biosensor consists of a pH-stable RFP fused with a pH-sensitive GFP. Under basal conditions, the mitochondrial network displays red and green fluorescence. In the context of mitophagy, as mitochondria undergo transportation to the acidic lysosomal environment, the GFP fluorescence undergoes quenching, while the RFP fluorescence remains stable [44]. The GFP/RFP ratio was quantified to assess mitophagy [20]. A heightened ratio, indicative of a decreased mitophagy level, was discerned in D8 and D12 worms compared to D4 worms. Interestingly, the 90-minute swim regimen induced mitophagy and recovery was evident 24 hours later in D4 and D8 worms, whereas D12 worms did not exhibit a change in mitophagy (Figure 3C). The obvious alterations of GFP/RFP during ageing and exercise were not observed in *prdx-2* mutant strain at any age (Figure 3C). These findings demonstrate that ageing results in an increase in mitochondrial fragmentation and decreased mitophagy. Exercise promotes mitochondrial fragmentation and mitophagy but the recovery was inhibited with age. The results highlight that loss of PRDX-2 results in increased mitochondrial fragmentation and potentially disrupted mitochondrial fusion along with a failure to activate mitophagy with exercise.

**Figure 3.**
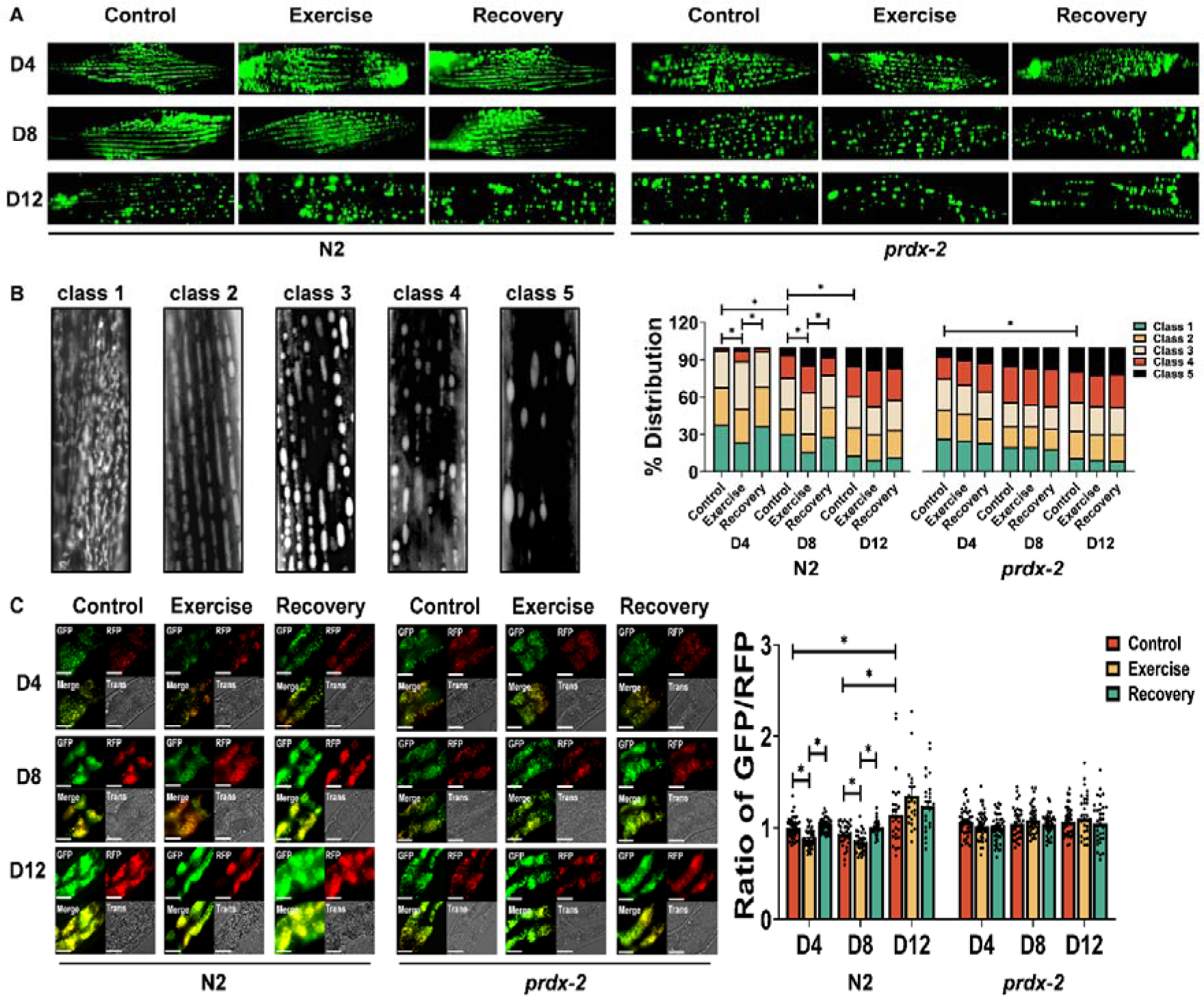
Ageing and loss of PRDX-2 results in mitochondrial fragmentation and disrupted mitophagy. (A-C) Representative images of *myo-3::gfp* reporter strain for mitochondrial morphology (A, B) and *myo3-p::tomm20::Rosella* reporter strain for mitophagy (C) following acute exercise at different ages, scale bar = 50 μm, n = 130-150 for mitochondria reporter, n = 30-45 for mitophagy reporter. Graphs are the normalised relative means ± SEM and all experiments were performed and *p*-value of < 0.05 was considered as statistically significant *(*p* < 0.05). *p* values (B: in N2 worms: D4: C vs E = 0.03, E vs R = 0.0456; D8: C vs E = 0.041, E vs R = 0.0478; D4 vs D8 = 0.0004; D8 vs D12 = 0.0007; in *prdx-2* worms: D4 vs D12 = 0.0135. C: in N2 worms: C vs E < 0.0001, E vs R < 0.0001; D8: C vs E = 0.0063; E vs R < 0.0001; D4 vs D12 = 0.0194; D8 vs D12 = 0.0004)

### 2.4 Exercise induces DAF-16 nuclear localisation and promotes an acute increase in genes regulating mitochondrial morphology that is absent in the *prdx-2* mutant strain

To elucidate the molecular consequences of alterations in mitochondrial adaptation during exercise and ageing process, the *prdx-2* mutant strain was crossed with DAF-16 reporter strains OH16024 *daf-16(ot971[daf-16::GFP]) I* and TJ356 *zIs356 [daf-16p::daf-16a/b::GFP + rol-6(su1006)]*. Moreover, mRNA levels of genes associated with mitochondrial morphology were quantified in both the N2 and *prdx-2* mutant strains. The distribution of DAF-16::GFP was quantified within the body-wall muscle of *C. elegans*, specified into three categories: cytosolic, intermediate and nuclear (Figure 4A). The acute exercise induced DAF-16 nuclear localisation in D4 and D8 worms, but not in D12 worms (Figures 4A and S3A). The *prdx-2* mutant did not induce DAF-16 nuclear localisation following exercise at any stage. Previous studies have demonstrated that *prdx-2* mutants at L4 stage have elevated resistance to sodium arsenite, as a result of increased DAF-16 nuclear localisation [45]. We confirmed these results at the L4 stage but at D4 there was decreased resistance to sodium arsenite and DAF-16 nuclear localisation (Figures S3B and S3C). Inhibition of *spg-7* and *ppgn-1* expression by DAF-16 nuclear localisation, alleviates the negative regulation of mitochondrial fusion protein EAT-3 by mitochondrial proteases SPG-7 and PPGN-1 [18]. There was decreased expression of *ppgn-1* and increased expression of *eat-3* in N2 at adult D4 and adult D8 following exercise, but not in the *prdx-2* mutant (Figures 4B-4D). Furthermore, the expression levels of *dct-1* (ortholog of BNIP3 mitophagy receptor), *cpt-1* (ortholog of CPT1, a marker of long chain fatty acid oxidation and regulator of mitochondrial morphology) and *skn-1* exhibited a notable increase either immediately or 24 hours post-exercise, particularly in D4 worms (Figures 4E-4G). In contrast, D12 worms did not display a similar elevation in the expression of any of these genes following exercise. Notably, the *prdx-2* mutant strain did not demonstrate an observable upregulation of these genes (*dct-1*, *cpt-1* and *skn-1*) following exercise at any age (Figures 4E-4G). Moreover, it was observed that the expression levels of *eat-3* and *skn-1* exhibit a gradual increase with advancing age (Figures S3D and S3I). The results demonstrate that exercise induces DAF-16 nuclear localisation and expression of genes promoting mitochondrial remodelling, these changes are absent in old worms and the *prdx-2* mutant strain.

**Figure 4.**
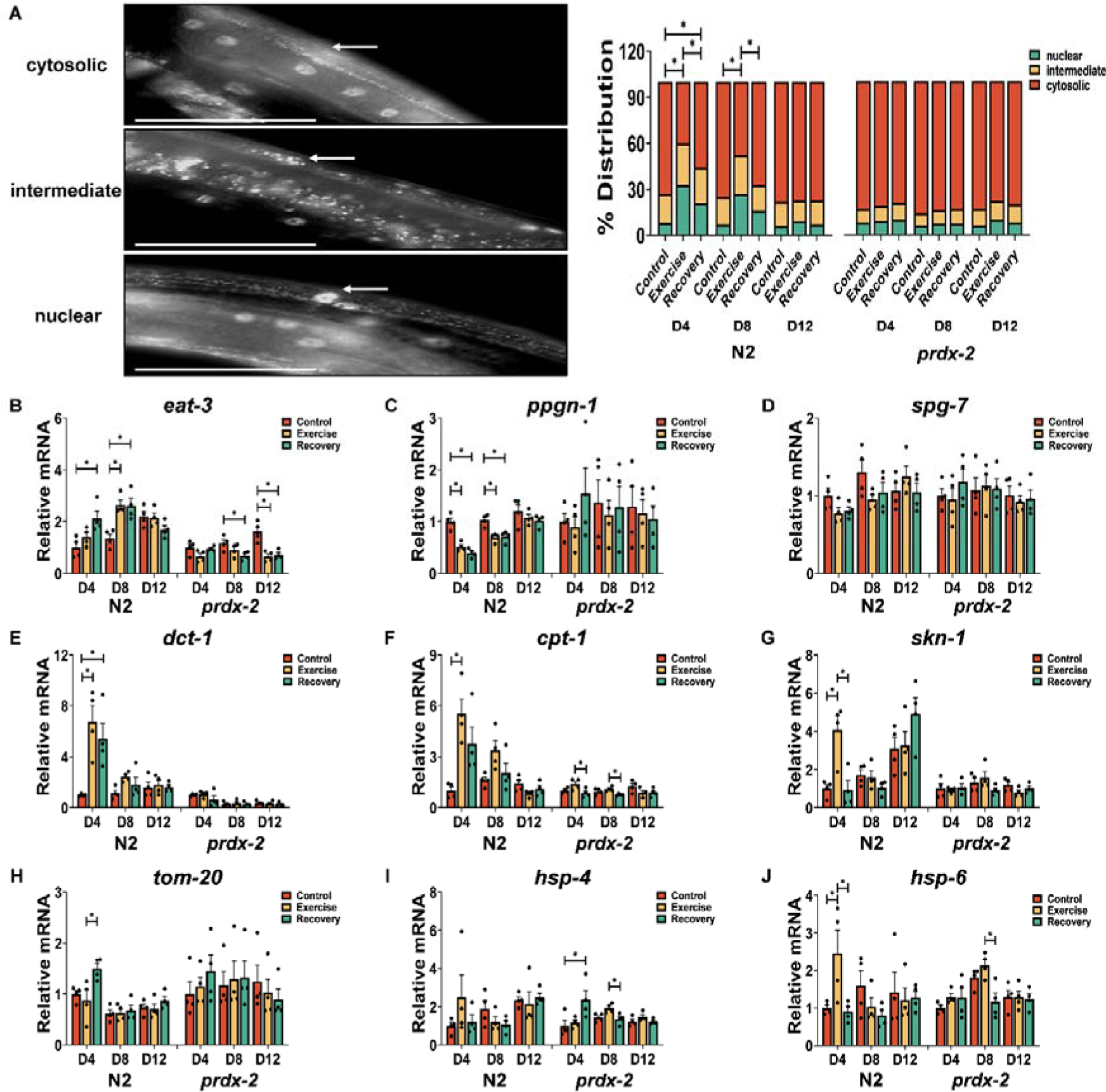
Ageing and loss of PRDX-2 adversely impacts the DAF-16 nuclear localisation and the expression of genes regulating mitochondrial dynamics and UPR in response to exercise. (A) Representative images of the OH16024 strain DAF-16::GFP distribution, scale bar = 50 μm, n= 130-150. (B-J) mRNA level of *eat-3* (B), *ppgn-1* (C), *spg-1* (D), *dct-1* (E), *cpt-1* (F), *skn-1* (G), *tom-20* (H), *hsp-4* (I) and *hsp-6* (J) following acute exercise at different stages, n = 4. Graphs are the normalised relative means ± SEM and all experiments were performed with n = 130-150 for DAF-16 reporter, n = 4 for mRNA expression and *p*-value of < 0.05 was considered as statistically significant *(*p* < 0.05). *p* values (A: in N2 worms: D4: C vs E < 0.0001, E vs R = 0.046, C vs R = 0.0182; D8: C vs E < 0.0001, E vs R = 0.0238. B: in N2 worms: D4: C vs R = 0.0157, D8: C vs E = 0.0059, C vs R = 0.0066; in *prdx-2* worms: D8: C vs R = 0.0409; D12: C vs E = 0.0019, C vs R = 0.0029. C: in N2 worms: D4: C vs E = 0.0004, C vs R < 0.0001; D8: C vs E = 0.0034, C vs R = 0.005. E: in N2 worms: D4: C vs E = 0.0073, C vs R = 0.0305. F: in N2 worms: D4: C vs E = 0.0063. G: in N2 worms: D4: C vs E = 0.0062, E vs R = 0.0055. H: in N2 worms: D4: E vs R = 0.0242. I: in *prdx-2* worms: D4: C vs R = 0.0467; D8: E vs R =0.0258. J: in N2 worms: D4: C vs E = 0.0449, E vs R = 0.0337; in *prdx-2* worms: D8: E vs R = 0.0108)

### 2.5 PRDX-2 is required for exercise induced UPR activation and mitochondrial ER remodelling

Mitochondria and the ER are essential regulatory centres in cellular homeostasis, engaging in a synergistic relationship. Communication between these organelles is facilitated by MERCS, which have been identified as key regulators of both mitochondrial fission and fusion [21, 22, 24]. The activation of the adaptive Unfolded Protein Response (UPR) is linked with remodelling of ER and MERCS [23]. Expression of *hsp-4* (reflection of ER-associated UPR) and *hsp-6* (indication of mitochondrial UPR) was quantified following exercise and during the ageing process. Results indicated a heightened expression of *hsp-6* following exercise at D4 in the N2 strain (Figure 4J). Furthermore, the expression levels of *hsp-4* increased at D8 and D12 compared to D4 in the N2 strain (Figure S3K). Notably, the *prdx-2* mutant strain did not demonstrate an upregulation of *tom-20* and *hsp-6* following exercise at any age (Figures 4H and 4J). The link between activation of the UPR following ER stress has been suggested to regulate MERCS formation [46]. Transmission electron microscopy (TEM) imaging of mitochondria and ER was employed to quantify mitochondrial morphology and MERCS formation in response to the cellular adaptations to exercise [47]. In N2 worms, the acute exercise induces a reduction in mitochondrial length and aspect ratio (length/width), implying a transition in muscle mitochondrial morphology from rod-shaped to oval-shaped, but this alteration was not observed in *prdx-2* mutant strain (Figure 5A). Following 5 days of exercise, there was a notable increase observed in mitochondrial length, area and aspect ratio in the N2 strain (Figure 5B). These findings suggest an augmentation in mitochondrial mass as a consequence of the chronic exercise regimen. In the comparison between adult D4 and D8 worms, it was observed that *prdx-2* mutants exhibited a more fragmented mitochondria relative to N2 worms, consistent with the highly fragmented mitochondrial network in the *prdx-2* mutant strain (Figure 3A). Notably, both acute and chronic exercise interventions resulted in a significant reduction in the MERC distance, representing a closer coupling between mitochondria and ER in the N2 strain. Furthermore, a notable increase in the frequency of Endoplasmic Reticulum-Mitochondria Contact Compartments (ERMICCs: calculated as MERC length divided by the product of mitochondrial perimeter and MERC distance) [48] was observed following both acute and chronic exercise interventions in the N2 strain but not in the *prdx-2* mutant strain (Figures 5A and 5B). These findings substantiate the presence of a substantial and dynamic interplay between mitochondria and the ER in response to exercise. Together, these findings suggest that PRDX-2 plays a key role in regulating UPR activation and MERCS formation following exercise.

**Figure 5.**
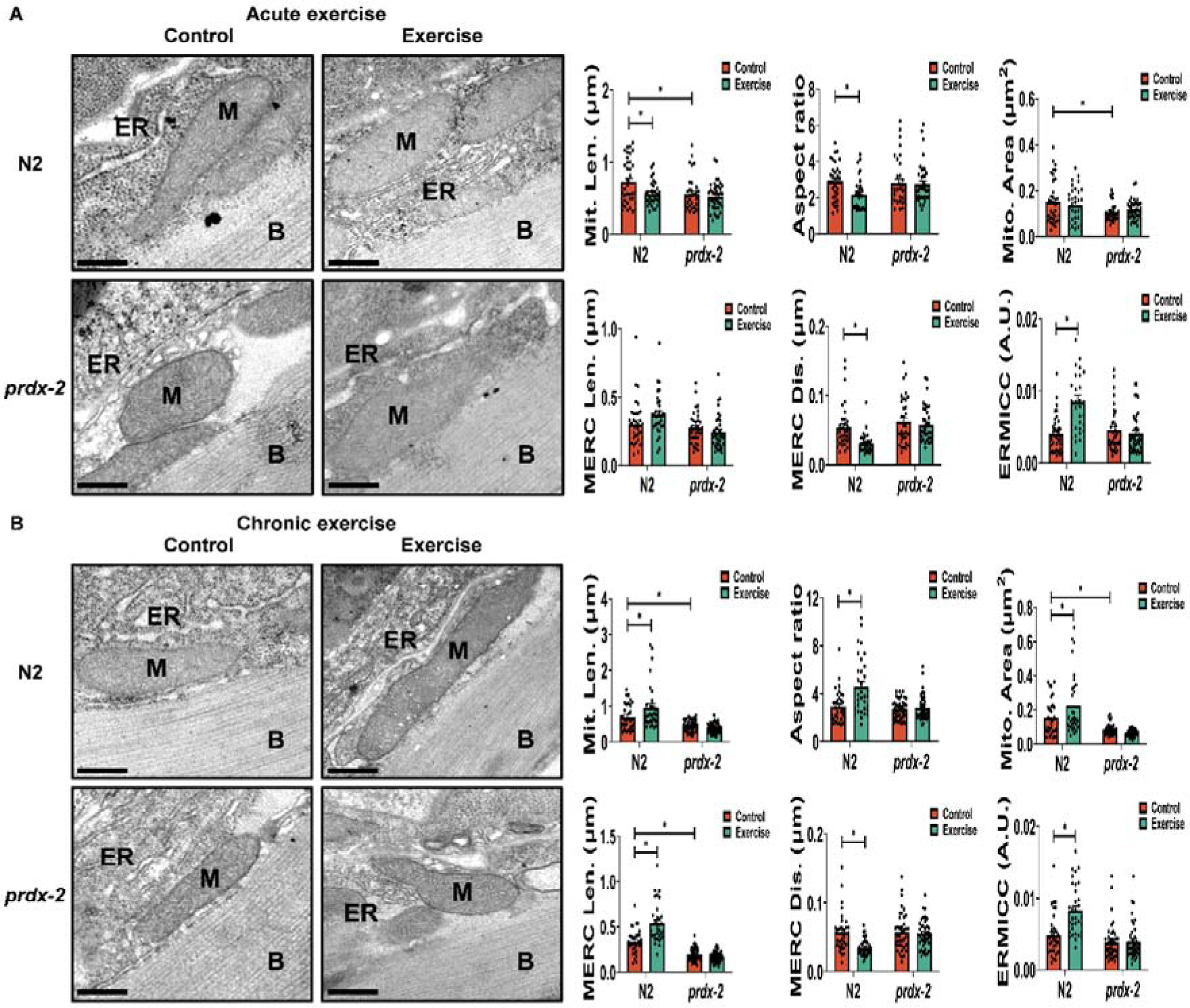
Exercise influences mitochondrial ER contact sites in N2 but not in the *prdx-2* mutant. (A-B) Mitochondria ER contact sites change following acute exercise (A) and chronic exercise (B), scale bar = 200 nm, n= 30-45. Graphs are the normalised relative means ± SEM and all experiments were performed and *p*-value of < 0.05 was considered as statistically significant *(p < 0.05). *p* values (A: mitochondrial length: N2: C vs E = 0.0231, N2 vs *prdx-2* = 0.0184. aspect ratio: N2: C vs E = 0.0317. mitochondrial area: N2 vs *prdx-2* = 0.0126. MERC distance: N2: C vs E = 0.0046. ERMICC: N2: C vs E < 0.0001. B: mitochondrial length: N2: C vs E = 0.0142; N2 vs *prdx-2* = 0.0162. aspect ratio: N2: C vs E < 0.0001. mitochondrial area: N2:C vs E = 0.0203; N2 vs *prdx-2* = 0.0225. MERC length: N2: C vs E < 0.0001; N2 vs *prdx-2* < 0.0001. MERC distance: N2:C vs E = 0.001. ERMICC: N2:C vs E < 0.0001)

### 2.6 PRDX-2 is required for increased fitness in response to exercise

To investigate the physiological functional implications of exercise and ageing in both N2 wild-type strain and *prdx-2* mutant strain that has an altered mitochondrial response capacity, CeleST [49] was implemented to quantify the activity patterns in these strains throughout the cycles of exercise and recovery at various stages of ageing. Parameters quantified included activity index, wave initiation rate, travel speed and brush stroke that are indicative of beneficial physical fitness, whereas body wave number, asymmetry, stretch and curling represent frailty [40]. In both strains, a consistent decline was observed in the activity index, travel speed, wave initiation rate and brush stroke with advancing age (Figures 6A, 6B, S4A and S4B). Simultaneously, there was an increase in body wave number, stretch, asymmetry and curling, exhibiting more pronounced effects in older worms, particularly at the D12 stage (Figures 6C, 6D, S4C and S4D). Moreover, acute exercise induced alterations in these parameters, reverting to baseline in D4 and D8 worms following exercise, but not in old D12 N2 worms. Importantly the *prdx-2* mutant strain did not recover following the exercise intervention at any age. In the comparative analysis of recovery rates in N2 at different ages, D4 worms exhibited superior recovery rates in travel speed, body wave number, stretch and wave initiation rate compared to D12 worms (Figures 6F-6H and S4E). Additionally, N2 demonstrated a more robust recovery rate in activity index, body wave number, stretch and wave initiation rate than *prdx-2* mutant, particularly notable at the D4 stage (Figures 6E, 6G, 6H and S4E). Furthermore, comprehensive quantification of these parameters over all stages revealed that the *prdx-2* mutant strain displayed impaired locomotory activity (Figure S4I). The data showed that physical fitness decreased with ageing and PRDX-2 is required for the enhanced locomotory activity or fitness in response to exercise.

**Figure 6.**
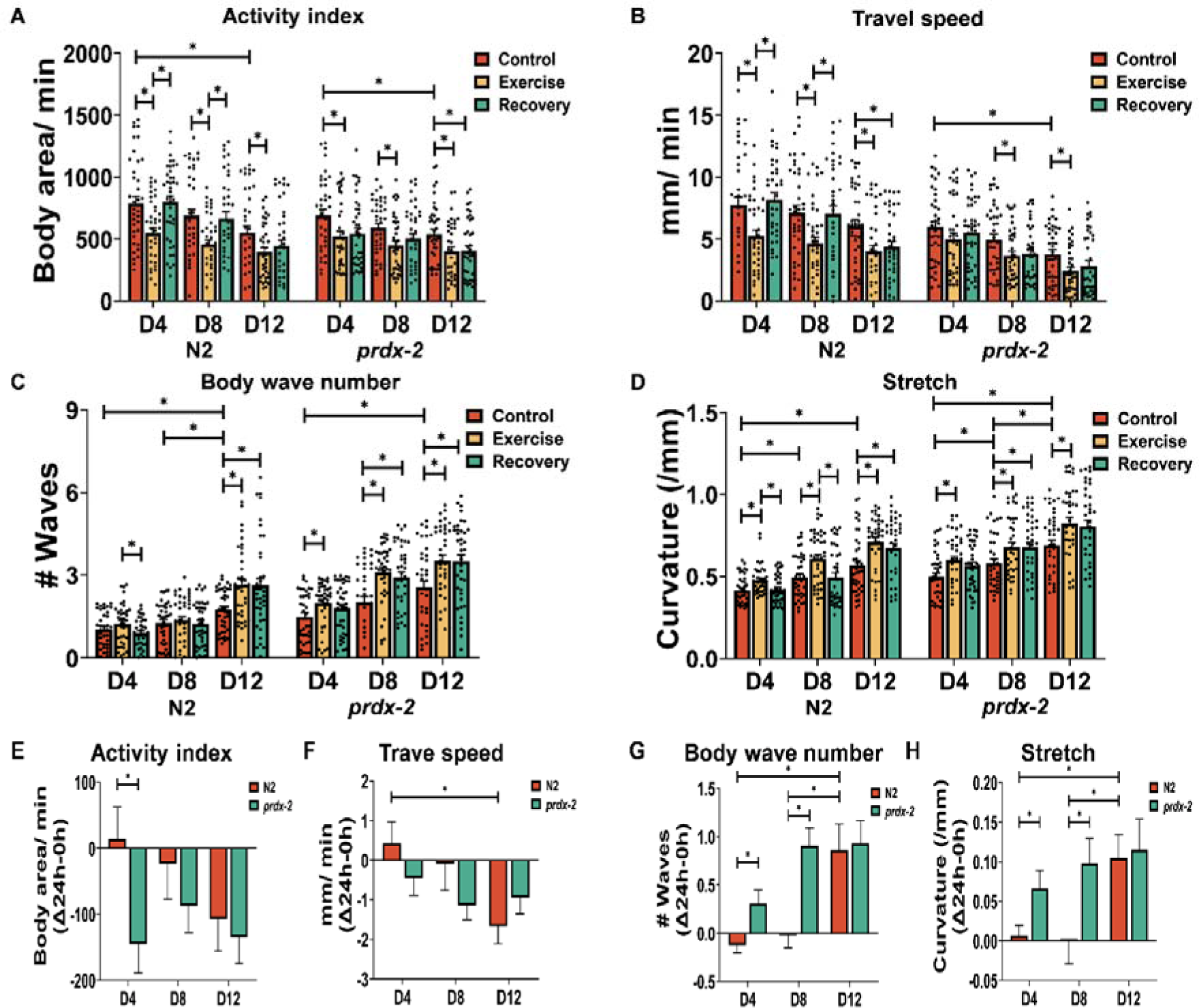
Ageing results in a decline in physical fitness and *prdx-2* mutants cannot recover following an exercise intervention. (A-D) Activity index (A), travel speed (B), body wave number (C) and stretch (D) following acute exercise at different stages, n = 30-40. (E-H) The recovery rate of activity index (E), travel speed (F), body wave number (G) and stretch (H) compared to normal condition, n= 30-40. Graphs are the normalised relative means ± SEM and *p*-value of < 0.05 was considered as statistically significant *(*p* < 0.05). *p* values (A: in N2 worms: D4: C vs E = 0.0032, E vs R = 0.0017; D8: C vs E = 0.0027, E vs R = 0.0088; D12: C vs E = 0.0405; D4 vs D12 = 0.0073; in *prdx-2* worms: D4: C vs E = 0.0262; D8: C vs E = 0.0231; D12: C vs E = 0.0291, C vs R = 0.0362; D4 vs D12 = 0.0289. B: in N2 worms: D4: C vs E = 0.006, E vs R = 0.001; D8: C vs E = 0.0098, E vs R = 0.0148; D12: C vs E = 0.008, C vs R = 0.0346. in *prdx-2* worms: D8: C vs E = 0.0441; D12: C vs E = 0.0263; D4 vs D12 = 0.0008. C: in N2 worms: D4: E vs R = 0.0365; D12: C vs E = 0.0084; C vs R = 0.0097; D4 vs D12 < 0.0001; D8 vs D12 = 0.0058; in *prdx-2* worms: D4: C vs E = 0.0394; D8: C vs E = 0.0002, C vs R = 0.0022; D12: C vs E = 0.0091, C vs R = 0.01; D4 vs D12 = 0.0003. D: in N2 worms: D4: C vs E = 0.0067, E vs R = 0.0177; D8: C vs E = 0.0064, E vs R = 0.0069; D12: C vs E = 0.0016, C vs R = 0.0267; D4 vs D8 = 0.036; D4 vs D12 < 0.0001; in *prdx-2* worms: D4: C vs E = 0.0098; D8: C vs E = 0.0467, C vs R = 0.0479; D12: C vs E = 0.0279; D4 vs D12 < 0.0001; D8 vs D12 = 0.0153; E: N2 vs *prdx-2*: D4 = 0.0165. F: in N2 worms: D4 vs D12 = 0.0217. G: in N2 worms: D4 vs D12 = 0.0003; D8 vs D12 = 0.0018; N2 vs *prdx-2*: D4 = 0.009; D8 = 0.0001. H: in N2 worms: D4 vs D12 = 0.0151; D8 vs D12 = 0.0095; N2 vs *prdx-2*: D4 = 0.0268; D8 = 0.00221)

In summary, the data presented demonstrate the adaptive rapid mitochondrial remodelling observed following acute exercise and ageing. PRDX-2 is sensitive to changes in the redox environment following exercise and required for exercise induced mitochondrial adaptations. Employing a cycle of acute exercise and recovery period at different stages in *C. elegans*, we observed that acute exercise initially induced mitochondrial fragmentation and mitophagy followed by mitochondrial fusion in young animals, but these responses decreased with age ultimately influencing physical fitness. Notably, the cycle of exercise and recovery failed to elicit these changes in the *prdx-2* mutant strain, evidenced by the failure to restore the redox environment, induce DAF-16 nuclear localisation and mitochondrial fusion, resulting in decreased physiological activity following exercise. Mechanistically, our results demonstrate that PRDX-2 is required for the nuclear localisation of DAF-16 following exercise and subsequent changes in mitochondrial morphology. Collectively, our data identify the indispensable role of PRDX-2 in orchestrating mitochondrial dynamics in response to a physiological stress by regulating DAF-16.

## 3 Discussion

In this study, we employed the nematode *C. elegans* as a model to explore the mechanistic role of PRDX-2 in regulating mitochondrial dynamics during exercise and ageing. Our data demonstrate that exercise induced alterations in the redox environment, resulting in increased DAF-16 nuclear localisation and MERCS formation dependent on PRDX-2, that regulates mitochondrial remodelling and physiological activity. These signalling mechanisms are disrupted in ageing and with the loss of PRDX-2, where there is a failure to restore the redox environment and results in reduced mitochondrial fusion and increased mitochondrial fragmentation due to an inability to stimulate DAF-16 nuclear localisation.

Mitochondrial quality is intricately regulated by mitochondrial biogenesis and selective degradation, which collectively contribute to the maintenance of mitochondrial mass, morphology and size [7, 50]. The dysregulation of mitochondrial biogenesis and impaired mitophagy are both recognised as contributing factors to compromised mitochondrial function in skeletal muscle during the ageing process [50–52]. Our previous studies have explored the effects of chronic swimming exercise on enhancing mitochondrial respiration, promoting mitochondrial biogenesis and regulating mitochondrial dynamics [20]. Recent research indicates that a cycle of mitochondrial fragmentation is induced during a single exercise session, followed by fusion after the recovery period [42]. This observation is corroborated by our current findings, which demonstrate that acute swimming triggers mitophagy, increases mitochondrial membrane potential and rapid changes in mitochondrial morphology. Furthermore, mitochondrial adaptations in response to exercise return to normal levels following a one-day recovery period. However, this cycle of mitochondrial remodelling was not observed in ageing worms as a result of an altered redox state of PRDX-2 and chronically elevated ROS stress resulting in disrupted DAF-16 nuclear localisation and downstream signalling.

A single swim session induces locomotory fatigue, elevated mitochondrial ROS and increased mitochondrial metabolic rate [41]. Our data demonstrates that exercise-induced ROS are associated with specific changes in the redox state of PRDX-2, increased DAF-16 nuclear localisation and mitochondrial remodelling that increases physical fitness and sensitivity to oxidative stress in young worms. However, these alterations were delayed or not observed in old worms. Recent studies have proposed a theory termed “redox-stress response capacity (RRC)” to elucidate this phenomenon, suggesting that cellular signalling and homeostasis maintenance are regulated through ROS-mediated adaptive responses or a hormesis effect [53]. Furthermore, the decline in RRC over time was termed as redox-stress response resistance (RRR), which offers an explanation for the reduction in RRC observed during ageing [54]. Interestingly, the mechanisms of RRC and RRR are mediated by PRDX-2 and hyperoxidation of Peroxiredoxins [54]. Decreased H_2_O_2_ sensitivity of PRDX-2 and elevated PRDX-2 hyperoxidation have been observed in ageing human fibroblasts and *C. elegans* [54]. These findings align with our own observations, as evidenced by reversible changes in the redox state of PRDX-2 following exercise only in young worms and a corresponding increase in hyperoxidised Peroxiredoxins in old worms. Furthermore, our previous redox proteomic analysis indicated that PRDX-2 regulates the induction of a redox relay for cellular adaptive response to exercise [20].

Peroxiredoxin 2, outcompetes other proteins for activation with physiological concentrations of H_2_O_2_ due to its abundance and kinetic reactivity [55]. However, Peroxiredoxin 2 has also been explored as involved in a redox relay for the transfer of oxidative equivalents to target proteins via reversible modification of Cysteine residues on redox-sensitive proteins [20, 54, 56]. STAT3, NRF2, and FOXO (DAF-16) are key transcription factors involved in the regulation of mitochondrial function and muscle myogenesis [17, 20, 57]. Peroxiredoxin 2 has been identified to interact with STAT3, forming a redox relay in response to a short bolus of H_2_O_2_ treatment [58]. Peroxiredoxin 1 has also been implicated in binding and regulating the activity of FOXO3 [59]. Likewise, the nuclear localisation of DAF-16 can be induced by ROS, with evidence suggesting that this process is regulated by its redox-sensitive cysteine residues forming a mixed disulphide with a redox signalling cascade [16]. However, the precise relationship between PRDX-2 and DAF-16 remains ambiguous in the current understanding of cellular signalling pathways.

DAF-16 is the only ortholog of the FOXO transcription factors in *C. elegans* and vital for preserving cellular homeostasis, it is required for the lifespan extension reported in *daf-2* mutant strains [15]. FOXO1 and FOXO3 govern glucose metabolism, lipid homeostasis and mitochondrial function primarily through their nuclear accumulation [60]. Elevated levels of ROS, originating from three mitochondrial mutants (*clk-1*, *isp-1* and *nuo-6*), prompt the nuclear translocation of DAF-16, thereby enhancing longevity [17]. The oxidation of Cysteine residues within FOXO/DAF-16 is necessary for its nuclear localisation and subsequent activation in response to ROS [16]. A previous study demonstrated that DAF-16 exhibits increased nuclear localisation in response to elevated ROS levels as well as following a swimming protocol [17]. Moreover, it is noteworthy that the heightened nuclear localisation of DAF-16 promotes mitochondrial fusion by promoting EAT-3 expression through the inhibition of mitochondrial proteases *spg-7* and *ppgn-1* [18]. Our findings corroborate that acute exercise promotes DAF-16 nuclear accumulation, leading to an increase in mitochondrial dynamics. Furthermore, we observed increased levels of the mitophagy receptor *dct-1*, mitochondrial fusion regulator *eat-3* and decreased *ppgn-1* following acute exercise. Surprisingly, these changes were not observed in the *prdx-2* mutant strain, suggesting that PRDX-2 is required for maintaining mitochondrial quality via regulating DAF-16 in response to exercise.

The altered redox state of PRDX-2 following exercise can potentially trigger the downstream activation of SKN-1, which has been reported via the p38 MAPK cascade [61]. It is widely recognised that changes occur in the transcriptome, proteome and metabolome during the ageing process [9]. RNA sequencing data from isolated tissues of ageing *C. elegans* demonstrated elevated expression levels of *skn-1*, *daf-16* and *prdx-2* in body wall muscle [62]. Similarly, it was demonstrated elevated levels of *daf-16* and *skn-1* in adult D7 worms compared to adult D1 worms [63]. These findings are consistent with the results presented here, which demonstrate heightened SKN-1 reporter activity and increased PRDX-2 (including hyperoxidised) protein levels at D12 compared to D4 worms. The *prdx-2* mutant strain displays an accelerated ageing phenotype [20, 43], characterised by a highly fragmented mitochondrial network evident at D4, which is aggravated with age. These findings suggest a potential role for PRDX-2 in maintaining mitochondrial integrity during both acute exercise and the ageing process. The depletion of PRDX-2 has previously been reported to enhance the activation of DAF-16 in response to low concentrations of arsenite [45]. These experiments were performed at the L4 stage in the *prdx-2* mutant strain. The accelerated ageing phenotype characteristic of the *prdx-2* mutant strain and altered redox environment could have resulted in the increased nuclear localisation of DAF-16 observed at this early developmental age.

In summary, our results demonstrate that there is blunted mitochondrial remodelling with age which is associated with an altered redox environment. PRDX-2 is required for mitochondrial adaptations in response to exercise through the regulation of the intracellular redox environment and appropriate DAF-16 nuclear localisation. Our data demonstrate that during ageing, there are elevated levels of ROS, increased mitochondrial fragmentation, reduced survival capabilities and decreased locomotory activity. Employing a cycle of acute exercise and recovery period model at various stages in *C. elegans*, acute exercise induces mitochondrial fragmentation and subsequent mitochondrial fusion through the activation of DAF-16, ultimately influencing physical fitness. Notably in the *prdx-2* mutant strain, the cycle of exercise and recovery failed to induce the observed changes compared to the N2 strain. The *prdx-2* mutant strain had an absence of alterations in mitochondrial membrane potential, mitochondrial dynamics as well as an inability to resolve the altered redox environment and reestablish mitochondrial adaptation and physiological activity following exercise. The redox state of PRDX-2 is sensitive to the redox environment and decreased reversible modification of the dimer ratio of PRDX-2 during ageing correlated with decreased survival and physical fitness in *C. elegans*. There are several limitations to the work presented in this study. We acknowledge that mitochondrial mass and morphology could affect the uptake of MitoTracker Red and MitoSOX [64] used to estimate mitochondrial membrane potential and mitochondrial ROS generation both of which are altered as a result of ageing and exercise. Furthermore, a physical interaction between PRDX-2 and DAF-16 was not demonstrated and future research should encompass additional investigations for a comprehensive understanding of this relationship. Despite these limitations, this study demonstrates the pivotal role of PRDX-2 in regulating DAF-16 nuclear localisation and mitochondrial remodelling during exercise and ageing.

## 4. Reagents and Resources

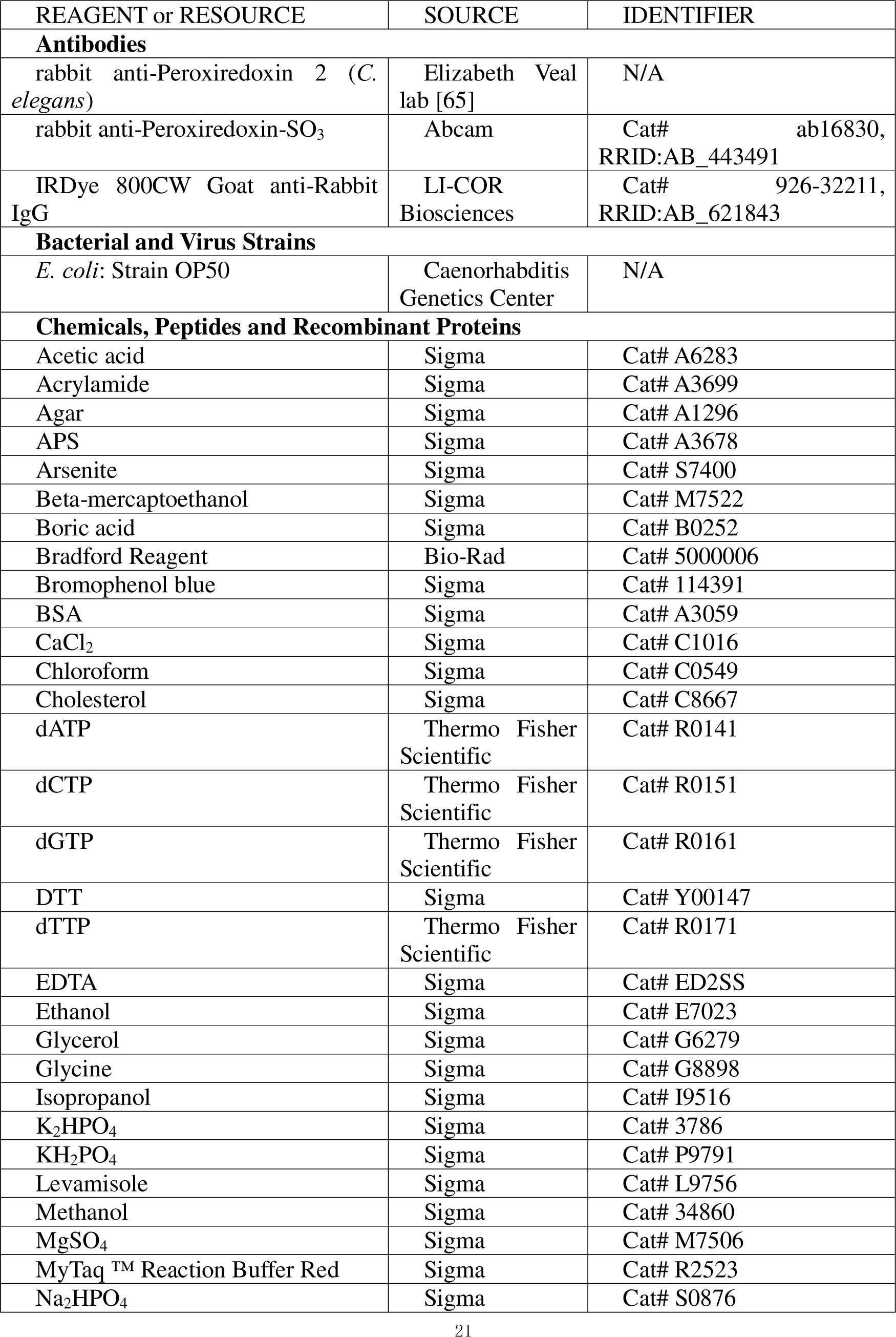
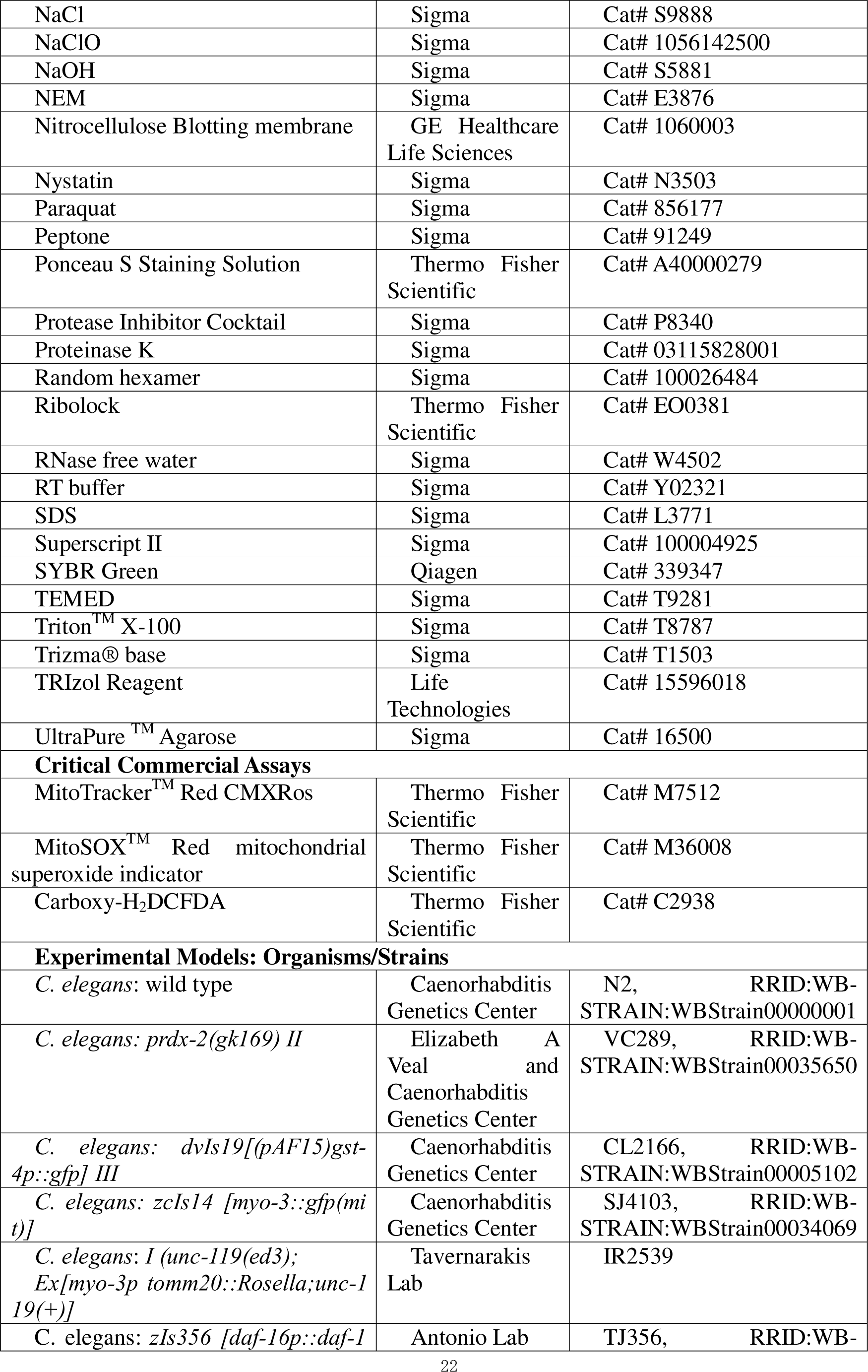
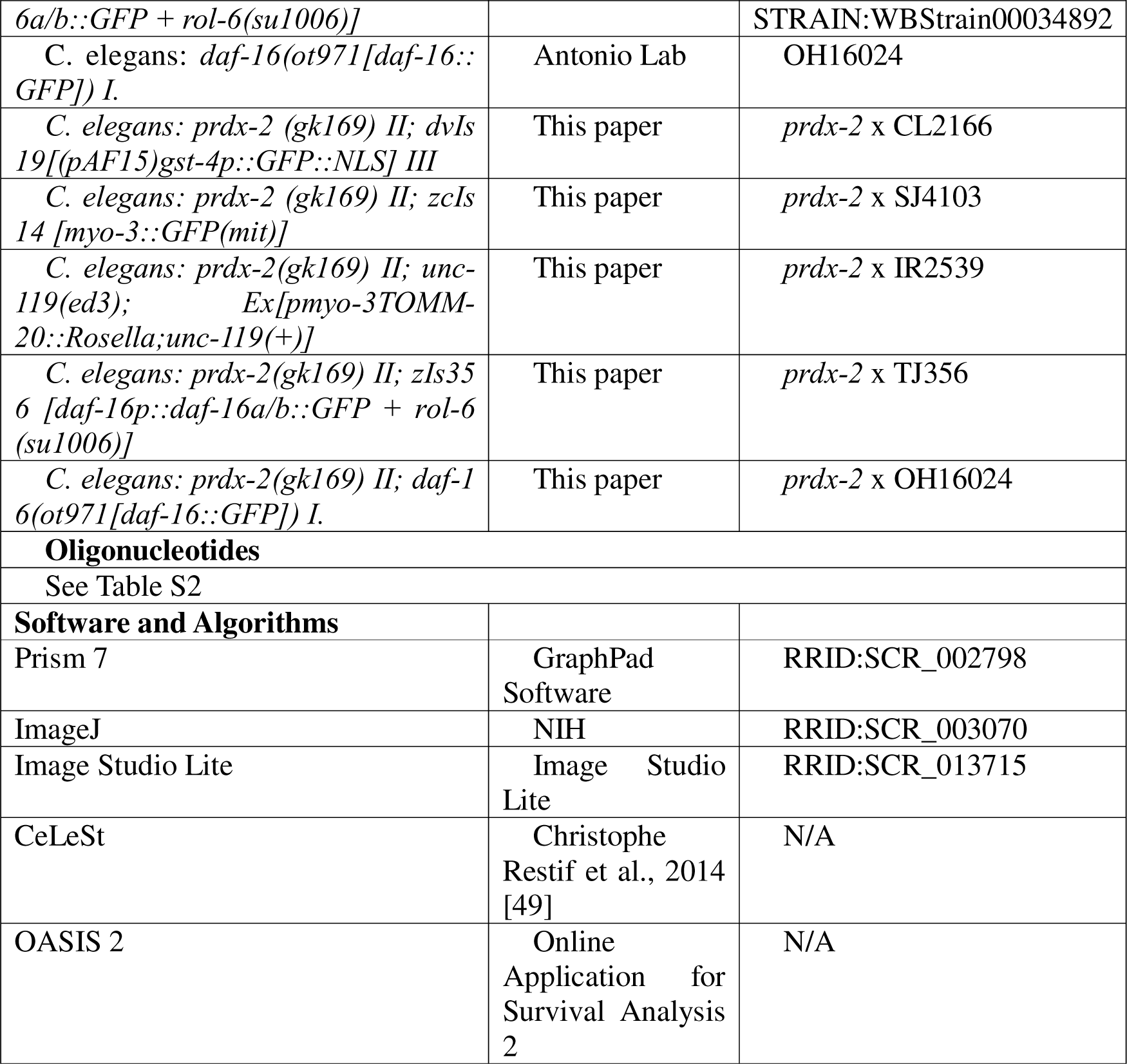

## 5. Materials and methods

### 5.1 C. elegans strains

*C. elegans* were cultured on NGM plates seeded with E. Coli (OP50) at 20 °C. *C. elegans* stains N2 wild type, VC289 (*prdx-2(gk169) II*) strain, CL2166 (*dvIs19[(pAF15)gst-4p::gfp] III)* and SJ4103 (*zcIs14 [myo-3::gfp(mit)]*) strains were obtained from the Caenorhabditis Genetics Center (CGC); IRE2539 (*Ex[pmyo-3 tomm-20::Rosella;unc-119(+)]*) was a gift from the Tavernarakis lab University of Crete, Greece; TJ356 (*zIs356 [daf-16p::daf-16a/b::GFP + rol-6(su1006)*) and OH16024 (*daf-16(ot971[daf-16::GFP]) I*) were obtained from Instituto de Biomedicina de Sevilla. In this paper, the *prdx-2* mutant strain was crossed with the reporter strains: SKN-1 reporter CL2166, DAF-16 reporter OH16024/TJ356, mitochondrial GFP reporter SJ4103 and mitophagy reporter IR2539.

### 5.2 C. elegans swimming exercise

For the different stages of worms, adult D1 worms were washed off plates and the larvae were removed daily to obtain worms at different ages (adult D4, D8, D12). For the acute exercise and control groups, 30 worms at least were transferred to unseeded NGM plates with or without M9 buffer, then worms were allowed to swim or crawl for a duration of 90 minutes at D4, D8 and D12 separately [41]. The recovery group underwent a 24-hour recovery period following the exercise regimen. For the chronic swimming exercise, adult D1 worms underwent 90-minute swimming sessions twice daily over a period of 5 days.

### 5.3 Oxidative stress survival assays

For the paraquat and sodium arsenite stress assays, 10 worms were selected for each group with a total of 60 worms per condition. These worms were carefully placed in individual wells of a 96-well plate containing either 100mM Paraquat or 2.5mM Sodium Arsenite, both diluted in M9 buffer. Worm death was determined by the absence of response to a gentle tap with a picker and survival was evaluated at intervals of 2 hours [66].

### 5.4 Western blotting

Protein samples from *C. elegans* were quantified using the Bradford reagent subsequent to homogenisation in an alkylating lysis buffer. 20 µg of protein per each condition was applied to 12% reducing or non-reducing SDS PAGE gels. The protein transfer was accomplished via a semi-dry blotter and the membrane underwent Ponceau S staining for normalisation. Following washing with TBS-T, the membranes were subjected to blocking in 5% milk within TBS-T for 1 hour at room temperature. Following the blocking step, the membranes were exposed to primary antibodies (rabbit anti-Peroxiredoxin 2 and rabbit anti-Peroxiredoxin-SO_3_) at a dilution of 1:1000 in 5% milk and this incubation persisted overnight. Subsequently, membranes were washed three times with TBS-T, each lasting 10 minutes. Membranes underwent incubation with the secondary antibody at a dilution of 1:10,000 in TBS-T, conducted in the absence of light, for a duration of 1 hour. Image acquisition was performed using the Odyssey Fc imaging system (Li-Cor). The subsequent analysis of blot quantification and normalisation was performed employing Image Studio Lite.

### 5.5 qPCR

RNA isolation from *C. elegans* was conducted utilizing TRIzol, following the methodology described in [67]. Subsequent to RNA isolation, cDNA synthesis and real-time qPCR were executed employing established protocols. For mRNA synthesis, 500 ng of RNA was mixed with 1 μl of random hexamers and incubated at 65 °C for 10 minutes. Then, the mixture was combined with a master mix including 4 μl of RT buffer, 2 μl of DTT, 1 μl of dNTP, 1 μl of Superscript II and 1 μl of Ribolock, followed by incubation at 42 °C for 60 minutes. Subsequent qPCR analysis was conducted using the SYBR Green Master Mix in a 10 µl reaction volume. The quantification of gene expression, relative to the housekeeping gene CDC42, was determined utilising the delta Ct method.

### 5.6 Imaging of C. elegans

For the staining of *C. elegans*, a total of 30-45 worms per condition were subjected to the procedure immediately after the acute exercise or following a 24-hour recovery period. The worms were incubated in 2.5 µM MitoTracker Red CMXRos for 10 minutes, 10 µM MitoSOX Red for 1 hour, or 25 µM H2DCFDA for 1 hour. After the probe incubation, the worms were transferred to NGM plates for 2-3 hours in the dark room to prevent accumulation of the stain in the guts. Subsequently, the worms were immobilised by 20 mM levamisole and imaged using EVOS at 10× magnification [20].

For the SKN-1 reporter, utilising the CL2166 (*dvIs19 [(pAF15)gst-4p::gfp III*) strain, a cohort of 30-45 worms at different ages underwent acute swimming. Post-exercise, the worms were immobilised on unseeded NGM plates for imaging using EVOS microscopy at a ×10 magnification. The assessment of green fluorescence intensity throughout the entire body of each worm was conducted using ImageJ [20].

For the assessment of mitochondrial dynamics in the body wall muscle, the SJ4103 strain (*zcIs14 [myo-3::gfp(mit)]*) was employed, using 30-45 worms subjected or not to acute swimming. Post-exercise, immobilisation of the worms was achieved using an agar slide covered with a coverslip. Subsequent imaging of mitochondria within the body wall muscles, specifically the region between the pharynx and vulva, was conducted under EVOS M7000 microscopy at a magnification of ×60. A total of 130-150 images were acquired and subsequently categorised into five distinct classes: Class 1, signifying highly abundant and well-networked mitochondria with minimal or no blebbing; Class 2, characterized by highly abundant mitochondria with network gaps and some blebbing; Class 3, featuring less abundant mitochondria with network gaps and increased blebbing; Class 4, indicating sparse mitochondria with some blebbing; and Class 5, denoting very sparse mitochondria [39].

To evaluate mitophagy, the IR2539 strain (*unc-119(ed3); Ex[pmyo-3 tomm-20::Rosella;unc-119(+)]*) was used. A group consisting of 30-45 worms underwent imaging at 60× magnification using the EVOS M7000 microscope. The ratio of green to red fluorescence intensity within a representative head region of each individual worm was subsequently determined using ImageJ [20].

To determine the nuclear localisation of DAF-16, the OH16024 strain (*daf-16(ot971[daf-16::GFP]) I.*) and TJ356 strain (*zIs356 [daf-16p::daf-16a/b::GFP + rol-6(su1006)]*) were used. 30-45 worms were immobilised using an agar slide covered with a coverslip. Subsequent imaging of DAF-16 nuclear localisation within the body wall muscles, specifically the region between the pharynx and vulva, was conducted under EVOS M7000 microscopy at a magnification of ×60. A total of 130-150 images were acquired and subsequently categorized into three distinct classes: nuclear, where the DAF-16::GFP distribution was visible in the nuclear of body wall muscle; intermediate, where the DAF-16::GFP distribution was not completely visible, showing punctate fluorescence in the cytosolic; and cytosolic, where the DAF-16::GFP distribution was in the cytosolic [18].

### 5.7 Transmission Electron Microscopy

1. *C. elegans* strains were immobilised with osmium tetroxide. Subsequently, the specimens underwent a dehydration process through a graded series of ethanol concentrations (30%, 50%, 70%, 90% and finally 100%), with each step lasting 2 × 15 minutes. Following the last 100% ethanol dehydration step, acetone was used to replace ethanol in a 2 × 20-minute procedure. Subsequently, the samples were immersed in a 50:50 mixture of resin and acetone for 4 hours, followed by placement in a 75:25 mixture on a rotator overnight. The following day, samples were transferred into 100% resin and rotated for 5-6 hours. Upon completion of this infiltration step, the specimens were relocated into the appropriate embedding mould, filled with fresh 100% resin and then subjected to an oven at 65 °C for 48 hours for polymerisation. Finally, the moulded samples were sectioned to prepare them for imaging. 10 captured images at least and 30-45 mitochondria were acquired at a magnification of 25,000 X. Using Image J according to [47], mitochondrial length (mitochondrial longitudinal distance), mitochondrial width (mitochondrial lateral distance), aspect ratio (ratio of mitochondrial length/ width), mitochondrial area (mitochondrial area size), MERC length (contacted length between mitochondria and ER), MERC distance (contacted distance between mitochondria and ER) and ERMICC (ratio of MERC length to product of mitochondrial perimeter and MERC distance),.

### 5.8 CeleST

To evaluate the swimming proficiency of *C. elegans*, CeleST analysis was performed following the acute exercise and the revery period. A minimum of 30 worms, arranged in 4-5 individuals per trial, were positioned on a glass substrate within a 10 mm ring. Subsequently, 60-second video recordings were captured at a rate of 16 frames per second, utilizing a Nikon LV-TV microscope set at 1× magnification and equipped with an OPTIKA C-P20CM camera [20, 40].

### 5.9 Statistical analysis

Images obtained from Western blot and microscopy of *C. elegans* were subjected to semi-automated quantification using ImageJ and Image Studio Lite and manual corrections were applied as needed to ensure accuracy in the quantification process. For the MitoTracker, MitoSOX, DCFDA and CL2166 strain staining, the quantification involved a total of 30-45 worms, with the fluorescence intensity measured per worm for each specific experimental condition. For SJ4103 strain staining, an evaluation was conducted on 130-150 images captured in the region between the pharynx and vulva and assessment was carried out based on the five predefined categories described earlier. For TJ356 and OH16024 strain, 130-150 images of body wall muscle were captured and quantified based on the DAF-16::GFP distribution categorized as nuclear, intermediate and cytosolic. For the IR2539 strain staining, the assessment involved determining the ratio of green fluorescence to red fluorescence for each individual worm. The specific details regarding the statistical analyses conducted are provided in the respective figure legend. To assess differences between two groups, a student t-test was employed. For comparisons involving more than two groups, either one-way or two-way ANOVA was utilised. The log-rank test was employed for comparing tolerance in the oxidative stress assay. The distribution into multiple classes was analysed using the chi-square test. A significance threshold of p-value < 0.05 was considered statistically significant. All graphical representations and statistical analyses were performed using GraphPad Prism 7. The source data used for figure generation in this manuscript are available in the source data file.

## Acknowledgements

We would like to sincerely thank Elizabeth Veal lab for providing the anti-PRDX-2 antibody and VC289 *prdx-2* mutant strain, the Tavernarakis lab for providing the IR2539 mitophagy reporter strain and the facilities and scientific and technical assistance of the Anatomy Imaging and Microscopy Facility at the University of Galway (https://imaging.universityofgalway.ie/imaging/) for electron microscopy work. QX (202006370047) and PL (202206370063) studentships are funded by the Chinese Scholarship Council (CSC), JCCM studentship is funded by the College of Nursing Medicine and Health Sciences, University of Galway.

## Author Contributions

Conceptualisation, BMcD and QX; methodology QX, PL, JCCM, KW, AMV, EM, PD and BMcD; investigation, QX, PLL, JCCM, EM and BMcD; resources, KW, AVM, PD and BMcD; writing-original draft QX and BMcD; writing-review and editing, QX, PL, JCCM, AMV, EM, PD, KW and BMcD; supervision, KW and BMcD.

---

**Figure S1.**
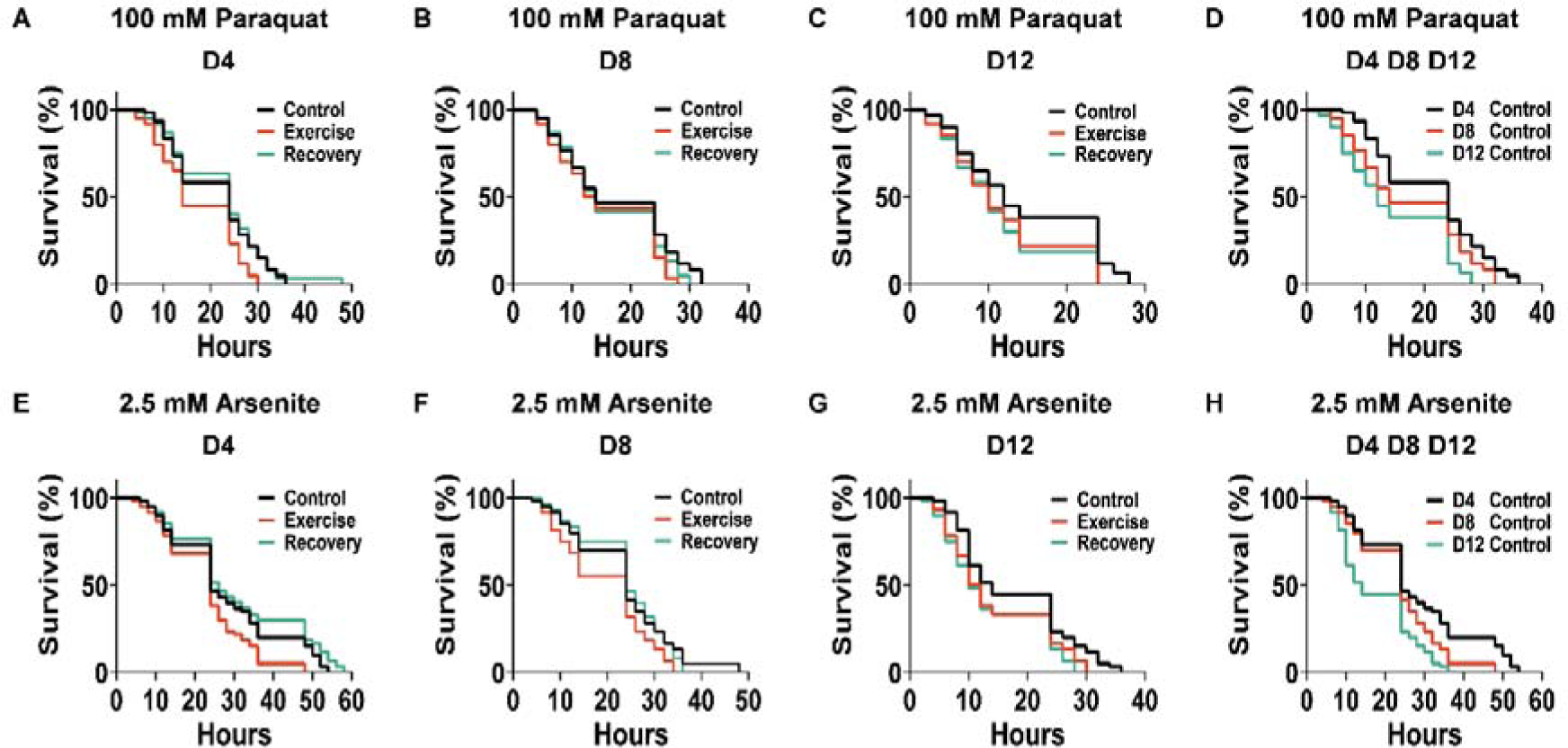
Decay of resistance to paraquat and arsenite with ageing and exercise. (A-D) Decreased survival to paraquat following acute exercise and ageing (A-D), n= 60. (E-H) Decreased survival to arsenite following acute exercise and ageing (E-H), n = 60. The Log-rank (Mantel-Cox) test was employed to compare survival between distinct groups. *p* values (A: C vs E *p*= 0.0034, E vs R *p*= 0.001; B: C vs E *p*= 0.0435; C: C vs E *p*= 0.022, C vs R *p*= 0.0076; D: D4 vs D8: *p*= 0.0266; D4 vs D12: *p* < 0.0001; D8 vs D12: *p*= 0.0181. E: C vs E *p*= 0.0114, E vs R *p*= 0.0009; F: C vs E *p*= 0.0143, E vs R *p*= 0.0079; G: C vs E *p*= 0.031; C vs R *p*= 0.0048; H: D4 vs D8: *p*= 0.0136; D4 vs D12: *p* < 0.0001; D8 vs D12: *p*= 0.0023)

**Figure S2.**
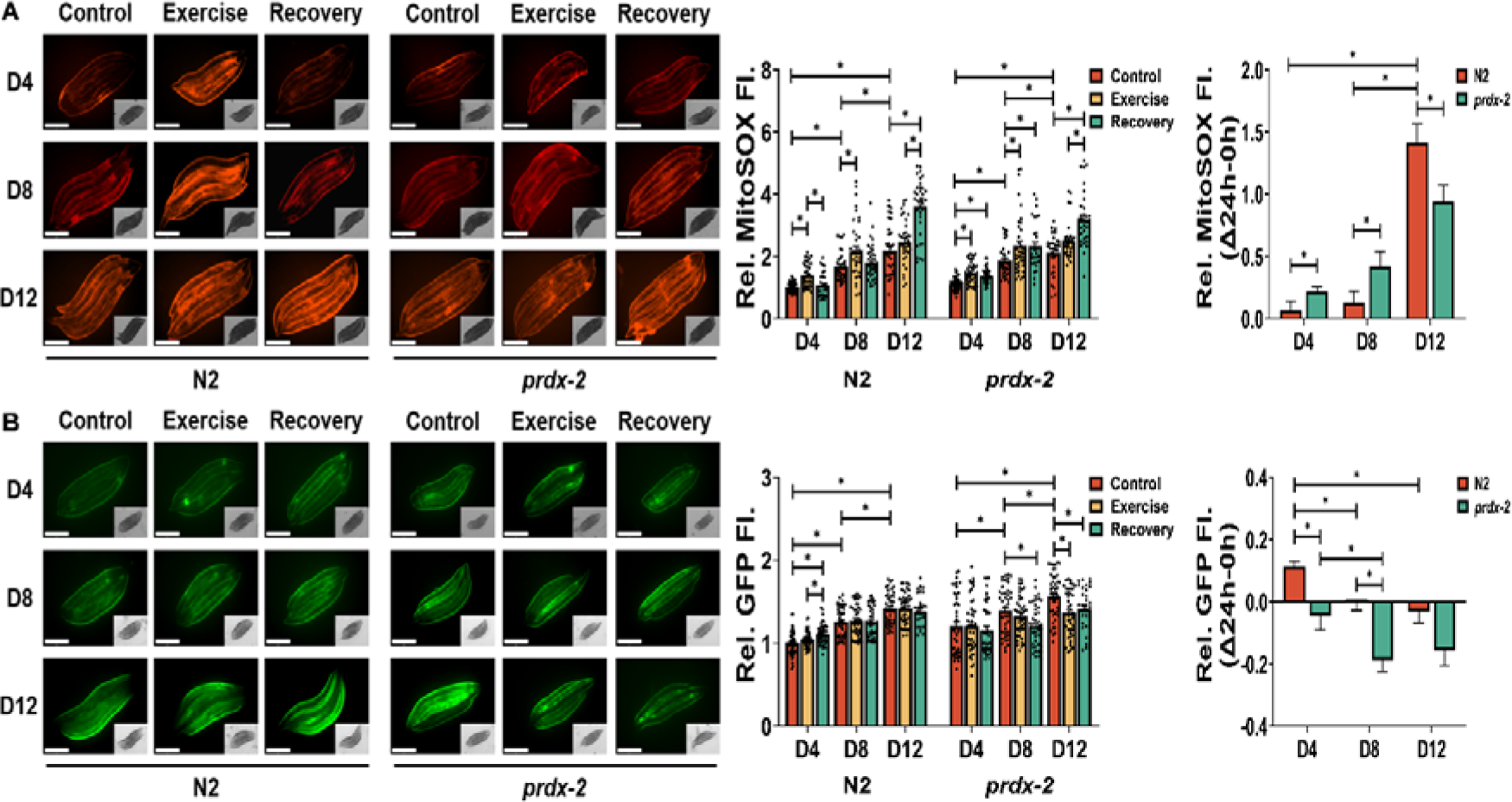
Ageing and the loss of PRDX-2 directly affect MitoSOX staining and SKN-1 activation. (A-B) MitoSOX red staining of worms for mitochondrial ROS (A) and *gst-4p::gfp* SKN-1 transcriptional reporter (B) following acute exercise at different stages, scale bar = 275 μm, n= 30-40. Graphs are the normalised relative means ± SEM and *p*-value of < 0.05 was considered as statistically significant *(*p* < 0.05), one-way or two-way ANOVA was used for significance between groups (a–b). *p* values (a: in N2 worms: D4: C vs E < 0.0001, E vs R = 0.0004; D8: C vs E = 0.0094; D12: C vs R < 0.0001, T vs R < 0.0001; D4 vs D8 < 0.0001; D4 vs D12 < 0.0001; D8 vs D12 = 0.0004; in *prdx-2* worms: D4: C vs E < 0.0001, C vs R = 0.0004; D8: C vs E = 0.0163, C vs R = 0.0315; D12: C vs T < 0.0001, E vs R = 0.0001; D4 vs D8 < 0.0001; D4 vs D12 < 0.0001; D8 vs D12 = 0.0332; recovery rate: N2: D4 vs D12 < 0.0001; D8 vs D12 < 0.0001; *prdx-2*: D4 vs D12 < 0.0001; D8 vs D12 = 0.0013; N2 vs *prdx-2*: D4 = 0.0453; D8 = 0.0486; D12 = 0.0211. b: in N2 worms: D4: C vs R < 0.0001, E vs R = 0.0217; D4 vs D8 < 0.0001; D4 vs D12 < 0.0001; D8 vs D12 < 0.0001. in *prdx-2* worms: D8: C vs R = 0.0024; D12: C vs E = 0.0014, C vs R = 0.045; D4 vs D8 = 0.0137; D4 vs D12 < 0.0001; D8 vs D12 = 0.0197. recovery rate: N2: D4 vs D8 = 0.0035; D4 vs D12 = 0.0007; *prdx-2*: D4 vs D8 = 0.0448; N2 VS *prdx-2*: D4 = 0.0018; D8 = 0.0001)

**Figure S3.**
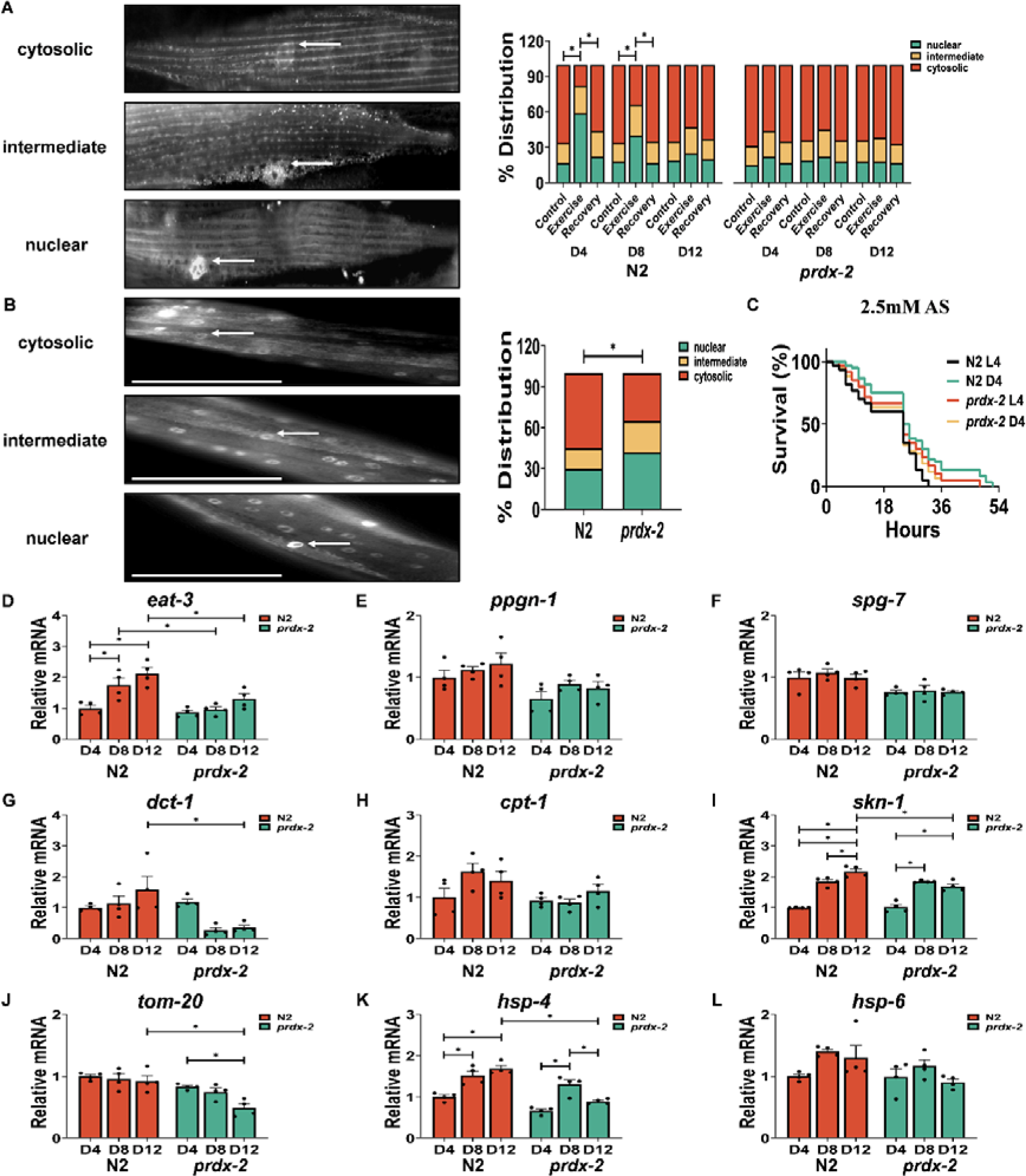
Ageing and loss of PRDX-2 in adult *C. elegans* adversely impacts DAF-16 nuclear localisation and mitochondrial dynamics in response to exercise. (A-B) Representative images of the TJ356 strain DAF-16::GFP distribution at adult (A) and L4 (B) stage in *C. elegans,* scale bar = 50 μm, n = 130-150. (C) Survival to arsenite in *prdx-2* mutant compared to N2 at L4 and D4 stage, n = 60. (D-L) mRNA level of *eat-3* (D), *ppgn-1* (E), *spg-7* (F), *dct-1* (G), *cpt-1* (H), *skn-1* (I), *tom-20* (J), *hsp-4* (K) and *hsp-6* (L) at different stages, n = 4. Graphs are the normalised relative means ± SEM and *p*-value of < 0.05 was considered as statistically significant *(*p* < 0.05). *p* values (A: in N2 worms: D4: C vs E < 0.0001, E vs R < 0.0001; D8: C vs E < 0.0001, E vs R < 0.0001. B: N2 vs *prdx-2* = 0.0172. C: N2 L4 vs *prdx-2* L4 = 0.0094; N2 D4 vs *prdx-2* D4 = 0.0186. D: in N2 worms: D4 vs D8 = 0.0274; D4 vs D12 = 0.0007. N2 vs *prdx-2*: D8 = 0.0188, D12 = 0.0135. G: N2 vs prdx-2: D12 = 0.0052. I: in N2 worms: D4 vs D8 < 0.0001, D4 vs D12 < 0.0001, D8 vs D12 = 0.0355; in *prdx-2*: D4 vs D8 < 0.0001, D4 vs D12 < 0.0001. N2 vs *prdx-2*: D12 = 0.0011. J: in *prdx-2* worms: D4 vs D12 = 0.0126; N2 vs *prdx-2*: D12 = 0.0013. K: in N2 worms: D4 vs D8 = 0.002, D4 vs D12 < 0.0001; in *prdx-2* worms: D4 vs D8 = 0.0002, D8 vs D12 = 0.0154; N2 vs *prdx-2*: D12: < 0.0001)

**Figure S4.**
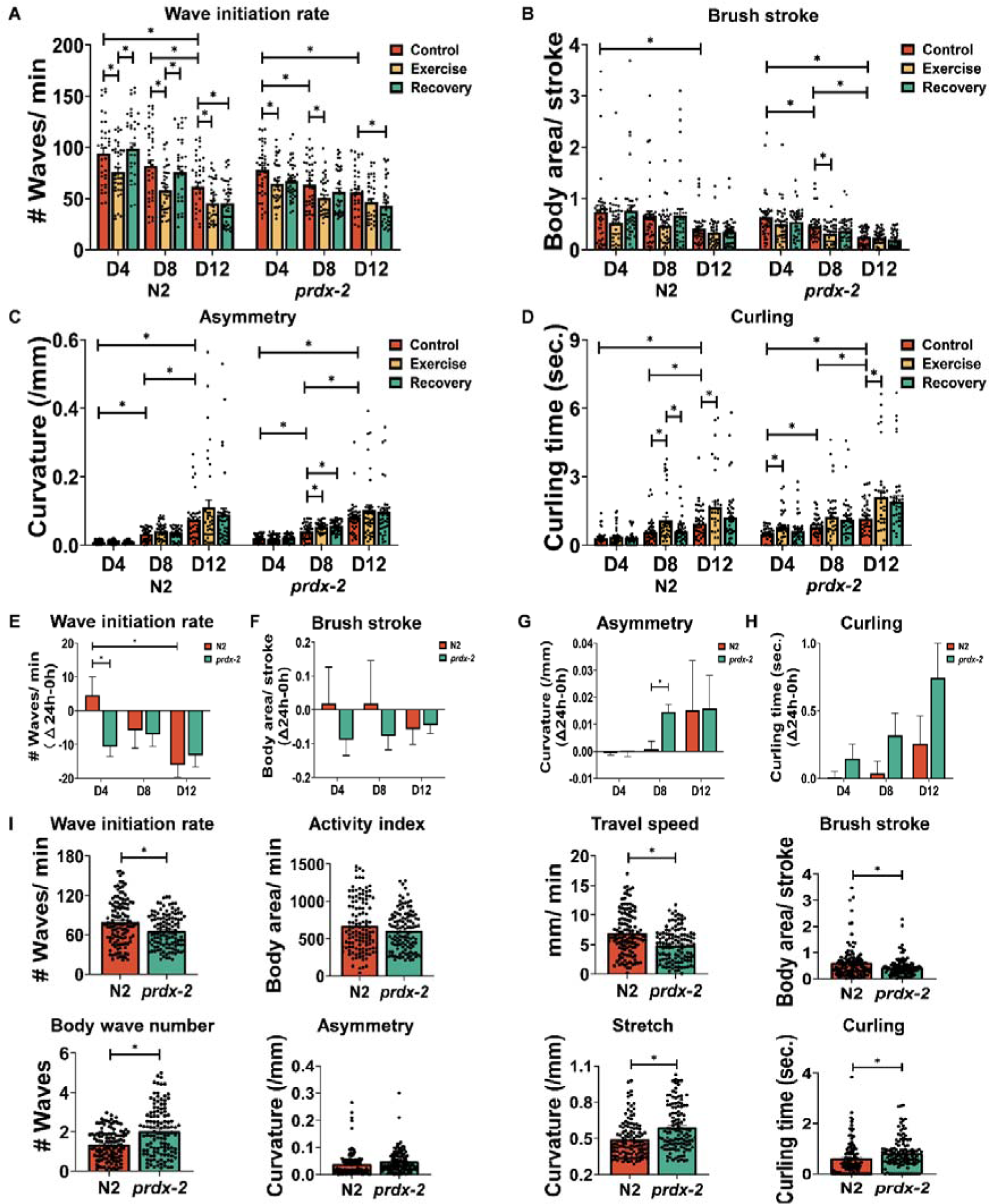
Ageing and loss of PRDX-2 undermines physical fitness following exercise. (A-D) Wave initiation rate (A), brush stroke (B), asymmetry (C) and curling (D) following acute exercise at different stages, n = 30-40. (E-H) The recovery rate of wave initiation rate (E), brush stroke (F), asymmetry (G) and curling (H) compared to normal condition, n = 30-40. (I) Physical fitness overall 12 days of N2 and *prdx-2* worms, n = 90-120. Graphs are the normalised relative means ± SEM and *p*-value of < 0.05 was considered as statistically significant *(*p* < 0.05). *p* values (A: in N2 worms: D4: C vs E = 0.027, E vs R = 0.0037; D8: C vs E = 0.0019, E vs R = 0.0278; D12: C vs E = 0.0063, C vs R = 0.0056; D4 vs D12 < 0.0001; D8 vs D12 = 0.014; in *prdx-2* worms: D4: C vs E = 0.0141; D8: C vs E = 0.022; D12: C vs R = 0.0179; D4 vs D8 = 0.0196; D4 vs D12 = 0.0001. B: in N2 worms: D4 vs D12 = 0.0433. in *prdx-2* worms: D8: C vs E = 0.0432; D4 vs D8 = 0.0318; D4 vs D12 < 0.0001; D8 vs D12 = 0.0133. C: in N2 worms: D4 vs D8 = 0.0164; D4 vs D12 < 0.0001; D8 vs D12 < 0.0001; in *prdx-2* worms: D8: C vs E = 0.016, C vs R = 0.0036; D4 vs D8 = 0.017; D4 vs D12 < 0.0001; D8 vs D12 < 0.0001. D: in N2 worms: D8: C vs E = 0.0161, E vs R = 0.0303; D12: C vs E = 0.0277; D4 vs D12 < 0.0001; D8 vs D12 = 0.013; in *prdx-2* worms: D4: C vs E = 0.0369; D12: C vs E = 0.014; D4 vs D8 = 0.0327; D4 vs D12 < 0.0001; D8 vs D12 = 0.0087. E: in N2 worms: D4 vs D12 = 0.0066; N2 vs *prdx-2*: D4 = 0.0126. G: N2 vs *prdx-2*: D8 = 0.001. I: wave initiation rate: N2 vs *prdx-2* = 0.0007; travel speed: N2 vs *prdx-2* < 0.0001; brush stroke: N2 vs *prdx-2* = 0.015; body wave number: N2 vs *prdx-2* < 0.0001; stretch: N2 vs *prdx-2* < 0.0001; curling: N2 vs *prdx-2* = 0.0098).

**Table S1.** Data from oxidative stress assays related to Figure 1B, 1C and S3C.

| Figures | age | Condition | n | Mean lifespan | Log-rank test, *p*-value |
| --- | --- | --- | --- | --- | --- |
| Figure 1B | D4 | Control | 60 | 21.1 ± 1.14 | |
| | | Exercise | 60 | 17.33 ± 1.05 | compared to D4 C, *p*= 0.0034 |
| | | Recovery | 60 | 21.83 ± 1.19 | compared to D4 E, *p*= 0.0011 |
| | D8 | Control | 60 | 17.57 ± 1.19 | compared to D4 C, *p*= 0.0266 |
| | | Exercise | 60 | 15.8 ± 1.08 | compared to D8 C, *p*= 0.0435 |
| | | Recovery | 60 | 16.5 ± 1.09 | compared to D8 E, *p*= 0.2379 |
| | D12 | Control | 60 | 14.7 ± 1.11 | compared to D8 C, *p*= 0.0181 |
| | | Exercise | 60 | 11.83 ± 0.93 | compared to D12 C, *p*= 0.022 |
| | | Recovery | 60 | 11.27 ± 0.89 | compared to D12 C, *p*= 0.0076 |
| Figure 1C | D4 | Control | 60 | 28.2 ± 1.97 | |
| | | Exercise | 60 | 23.37 ± 1.34 | compared to D4 C, *p*= 0.0114 |
| | | Recovery | 60 | 30.83 ± 2 | compared to D4 E, *p*= 0.0009 |
| | D8 | Control | 60 | 23.77 ± 1.34 | compared to D4 C, *p*= 0.0136 |
| | | Exercise | 60 | 19.67 ± 1.19 | compared to D8 C, *p*= 0.0143 |
| | | Recovery | 60 | 24.03 ± 1.12 | compared to D8 E, *p*= 0.0079 |
| | D12 | Control | 60 | 17.77 ± 1.21 | compared to D8 C, *p*= 0.0023 |
| | | Exercise | 60 | 14.6 ± 1.12 | compared to D12 C, *p*= 0.0231 |
| | | Recovery | 60 | 13.93 ± 1.08 | compared to D12 C, *p*= 0.0048 |
| Figure S3C | L4 | N2 | 60 | 19.3 ± 1.23 | |
| | | *prdx-2* | 60 | 22.9 ± 1.43 | compared to N2 L4, *p*= 0.0094 |
| | D4 | N2 | 60 | 26.47 ± 1.57 | |
| | | *prdx-2* | 60 | 21.77 ± 1.43 | compared to N2 D4, *p*= 0.0186 |

**Table S2.**
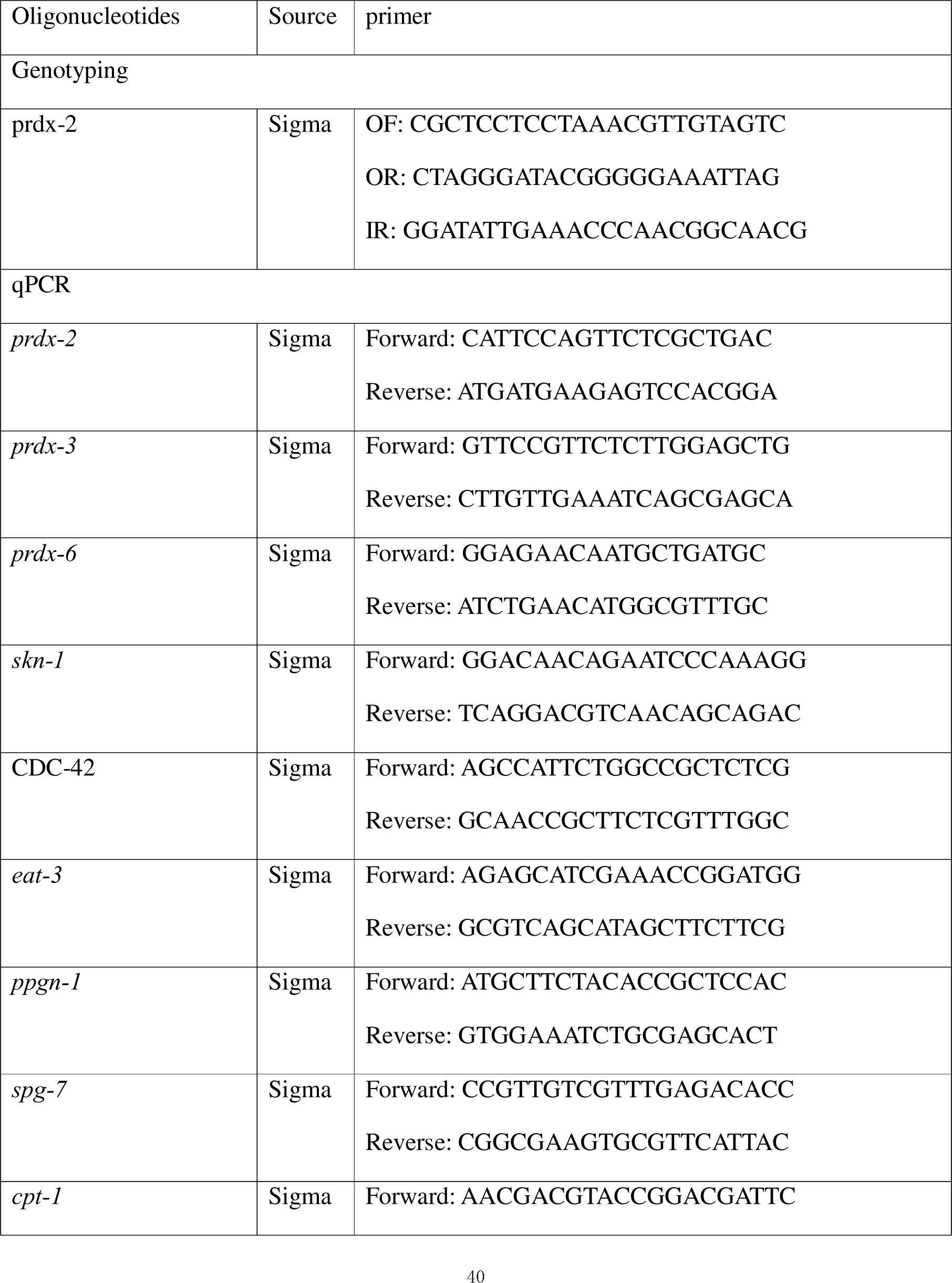
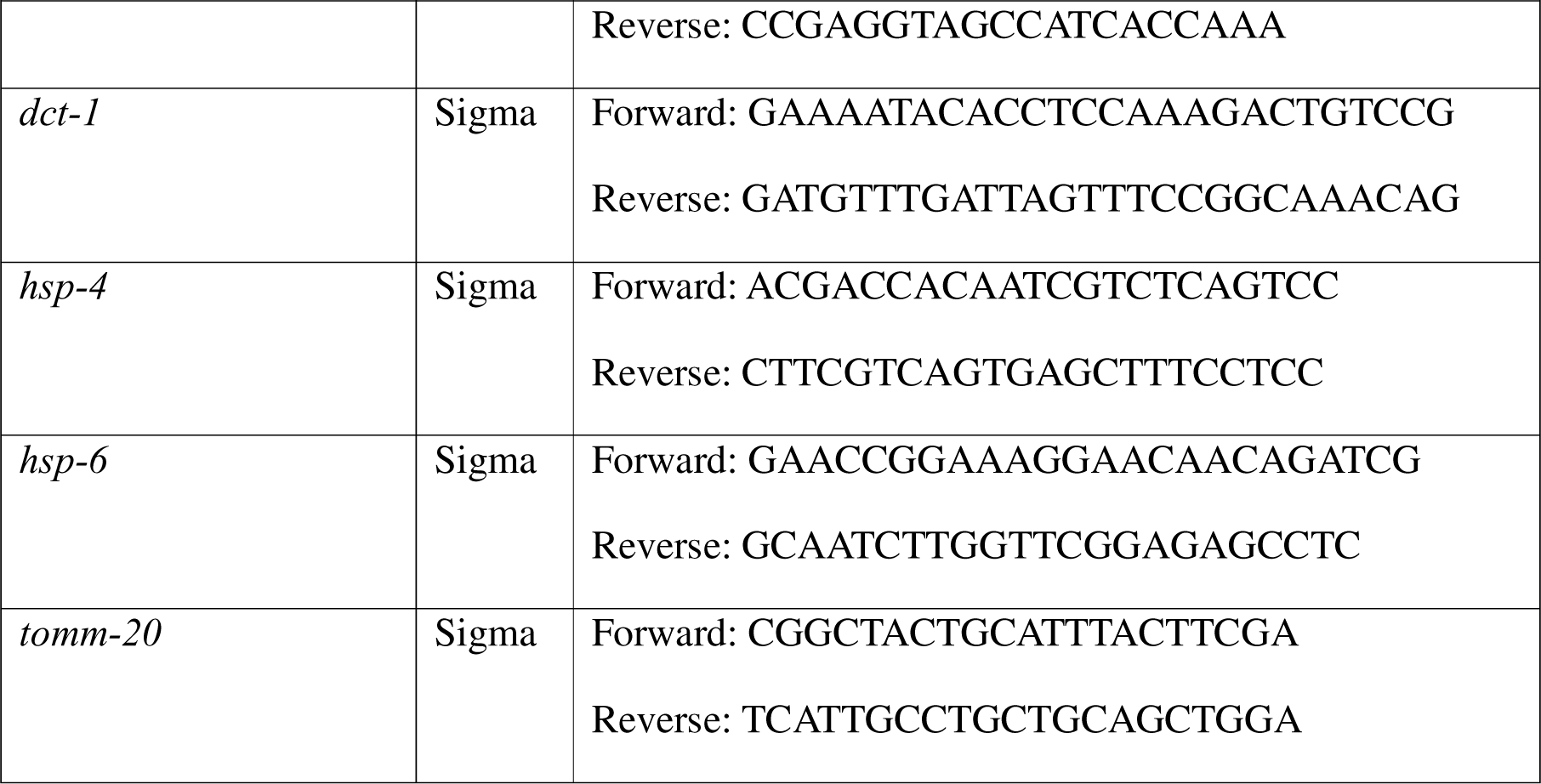
genotyping and qPCR primers.

